# Patient induced pluripotent stem cells identify specificities of a reticular pseudodrusen phenotype in age-related macular degeneration

**DOI:** 10.1101/2025.11.19.689393

**Authors:** Jenna C. Hall, Kavitha Krishna Sudhakar, Maciej Daniszewski, Anne Senabouth, Carla J. Abbott, Helena H. Liang, Himeesh Kumar, Grace E. Lidgerwood, Mehdi Mirzaei, Jessica Ma, Trevor Atkeson, Yumiko Hirokawa, Emeline F. Nandrot, Alexander Barnett, Chantal Cazevieille, Gaël Manes, Simon Mountford, Philip Thompson, Erica L. Fletcher, Zhichao Wu, Melanie Bahlo, Brendan R.E. Ansell, Daniel Paull, Alex W. Hewitt, Robyn H. Guymer, Joseph E. Powell, Alice Pébay

## Abstract

**Background:** Age-related macular degeneration (AMD) is a leading cause of vision loss. Reticular pseudodrusen (RPD), deposits on the apical side of the retinal pigment epithelium (RPE), signify a distinctive and critical AMD phenotype. Yet, their molecular basis and relationship to the conventional drusen seen in AMD remain unclear.

**Results:** We generated induced pluripotent stem cell-derived RPE cells from a clinically phenotyped cohort comprising only individuals with conventional drusen (AMD/RPD−) or drusen coexisting with RPD (AMD/RPD+). From these cells, we generated single-cell transcriptomics, proteomics, and functional data to identify differences between the two cohorts. We show that AMD/RPD+ RPE cells exhibit enrichment in extracellular matrix (ECM) remodelling, cytoskeletal, and hypoxia-responsive programs, whereas AMD/RPD− RPE cells display a relatively greater representation of mitochondrial and protein homeostasis pathways. Both subtypes engaged pathways classically linked to ageing, including ECM remodelling and mitochondrial function, but differed in the direction and extent of these changes. Expression and protein quantitative trait loci (QTLs) highlight shared genetic influences on mitochondrial and iron-handling pathways, while disease-interacting eQTLs and transcriptome-wide association study identify regulatory signals that are distinctive of the RPD subtype within AMD, including through regulation of ECM. Functionally, all iPSC-derived RPE formed drusen-like deposits *in vitro*: AMD/RPD−lines generated more basal deposits, whereas AMD/RPD+ cells exhibited greater structural instability under bisretinoid-induced stress.

**Conclusions:** These findings indicate that AMD with and without RPD represent mechanistically distinct entities and provide novel insight into the molecular mechanisms underlying disease heterogeneity in AMD.

## Background

Age-related macular degeneration (AMD) is a progressive, vision-threatening disease associated with dysfunction and death of the retinal pigment epithelium (RPE) and photoreceptors [1]. A pathological hallmark of AMD is accumulation of drusen deposits underneath the RPE [2]. The increasing size of drusen is significantly associated with a higher risk of progression to late-stage disease [3]. However, AMD patients can present with drusenoid deposits on the apical side of the RPE in the subretinal space, called reticular pseudodrusen (RPD) or subretinal drusenoid deposits [4]. The presence of RPD in patients with AMD has been reported to be a significant risk factor for progression to late-stage AMD [5–7], which includes geographic atrophy and choroidal neovascularisation [8,9]. The presence of RPD in individuals with AMD is also associated with significant visual function impairment [10,11], especially in dark adaptation [12,13]. Furthermore, there is evidence of treatment effect modification in the AMD/RPD+ phenotype in a randomized controlled clinical trial for AMD, suggesting different treatments may be required for these different phenotypes of AMD [14]. The molecular composition of RPD shares some similarities with that of conventional drusen; however, significant differences in lipid content have been observed [15,16], suggesting distinct pathways are involved in conventional drusen and RPD accumulation [17].

It is hypothesised that genetic and environmental factors participate in RPD development. Elucidation of contributing factors has been hindered by difficulties in detection and by inconsistent criteria used to identify RPD presence (number of lesions and type of imaging used), complicating comparisons between studies. Several studies have examined the relationship between AMD-associated genetic variants and the presence of RPD [18]. The *ARMS2* rs10490924 single nucleotide polymorphism (SNP) has been consistently reported as enriched in individuals with AMD and RPD compared to those AMD eyes without RPD [19–27]. Associations with the *CFH* rs1061170 SNP have also been described [19,21,23,24,26,27], although other studies failed to detect such a relationship [20,22,25]. Additional reported associations include variants in *C3* (rs22130199) [24], *C2/CFB* (rs641153) [24], *VEGFA* (rs943080) [24], *LIPC* (rs10468017) [27] and additional *CFH* SNPs (rs12144939, rs800292, rs393955, rs2274700) [23,24]. However, many earlier studies used colour fundus photography and incident-case designs, limiting reliable RPD identification and making it difficult to distinguish RPD−specific from general AMD genetic effects. As these studies largely focused on known AMD variants and lacked RPD−specific stratification [10], genome-wide approaches in well-characterised AMD cohorts are needed to clarify RPD−specific genetic risk factors. Recent genome-wide association studies (GWAS) have confirmed the association of *ARMS2/HTRA1* rs11200638 SNP with RPD and also identified new genetic variants associated with RPD, rs79641866/*PARD3B*, rs143184903/*ITPR1*, and rs76377757/*SLN* [28] and a long non-coding RNA gene *HTRA1-AS1* (ENSG00000285955/BX842242.1) [29].

The incidence of AMD increases with age, and many of the cellular processes implicated in its pathogenesis, including mitochondrial dysfunction, impaired protein homeostasis, extracellular matrix (ECM) remodelling, and chronic inflammation, mirror established hallmarks of aging [30]. Whether RPD reflects an accelerated form of these processes, or represents a distinct aging-associated trajectory remains unclear. As induced pluripotent stem cells (iPSCs) [31,32] can be differentiated into homogenous RPE cultures, and have been widely used for disease modelling, including for AMD [33–42], we modelled RPE biology *in vitro* using patient iPSC-derived RPE cells. These cells were from AMD donors, who were carefully phenotyped, and categorised as either with (AMD/RPD+) or without RPD (AMD/RPD−), focusing on their molecular programs, propensity to form drusen-like deposits, and resilience to stressors relevant to aging retina.

## Results

### Generation of iPSCs, differentiation into RPE cells, genomic profiling

Fibroblast cultures generated from skin biopsies of 103 individuals exhibiting either only drusen (AMD/RPD−) or drusen coexisting with extensive RPD (AMD/RPD+) were reprogrammed into iPSCs (72 females and 31 males; mean ± SD: 74.6 ± 7.9 years at recruitment) (**Figure S1**). The iPSC lines were genotyped for 845,487 SNPs and imputed with the Haplotype Reference Consortium panel [43]. Following quality control assessments, a total of 4,063,692 autosomal SNPs were obtained, with minor allele frequencies (MAF) >10%. To minimise experimental variation, all lines were differentiated into RPE cells in five independent batches as previously described [41], and suboptimally differentiated cultures (assessed by homogeneous cobblestone morphology and pigmentation) were discarded from analysis. This method yields functional RPE cells that express canonical RPE cell markers (**Figure S1**). Differentiated cell lines were divided into 13 pools, each consisting of up to 8 cell lines from both groups. scRNA-seq was performed on all pools, with the targeted capture of 20,000 cells per pool and a sequencing depth of 30,000 reads per cell. The resulting single-cell transcriptome profiles underwent quality control and donor assignment to remove cells designated as doublets or from individual lines that had low number of assignment anomalies identified by CNV array (10 lines) and failed genotyping. By the end of quality control assessment, 130,524 cells from 68 individual lines remained: 35 lines from participants with conventional AMD only (AMD/RPD−, 66,882 cells, seven males, 28 females, mean ± SD age: 71.1 ± 8.5 years) and 33 lines from patients with RPD (AMD/RPD+, 63,642 cells, 14 males, 19 females, mean ± SD age: 79.7 ± 1.0 years).

The cells were distributed among 16 subpopulations which were detected in both subphenotypes at comparable frequencies (**Table 1**), and classified across subtypes of neuronal progenitors, RPE progenitors and RPE cells. Consistent with previous observations [41], transcriptomic differences among the 14 RPE subpopulations primarily reflect changes in maturity, rather than in cell identity (**Figure 1**). For downstream analyses, these RPE clusters were therefore aggregated. Genes associated with AMD were expressed across all subpopulations in both cohorts (**Figure S2**). Of note, no significant difference in complement factor H (*CFH*) and *C3* were observed between cohorts (**Figure S2**).

**Figure 1.**
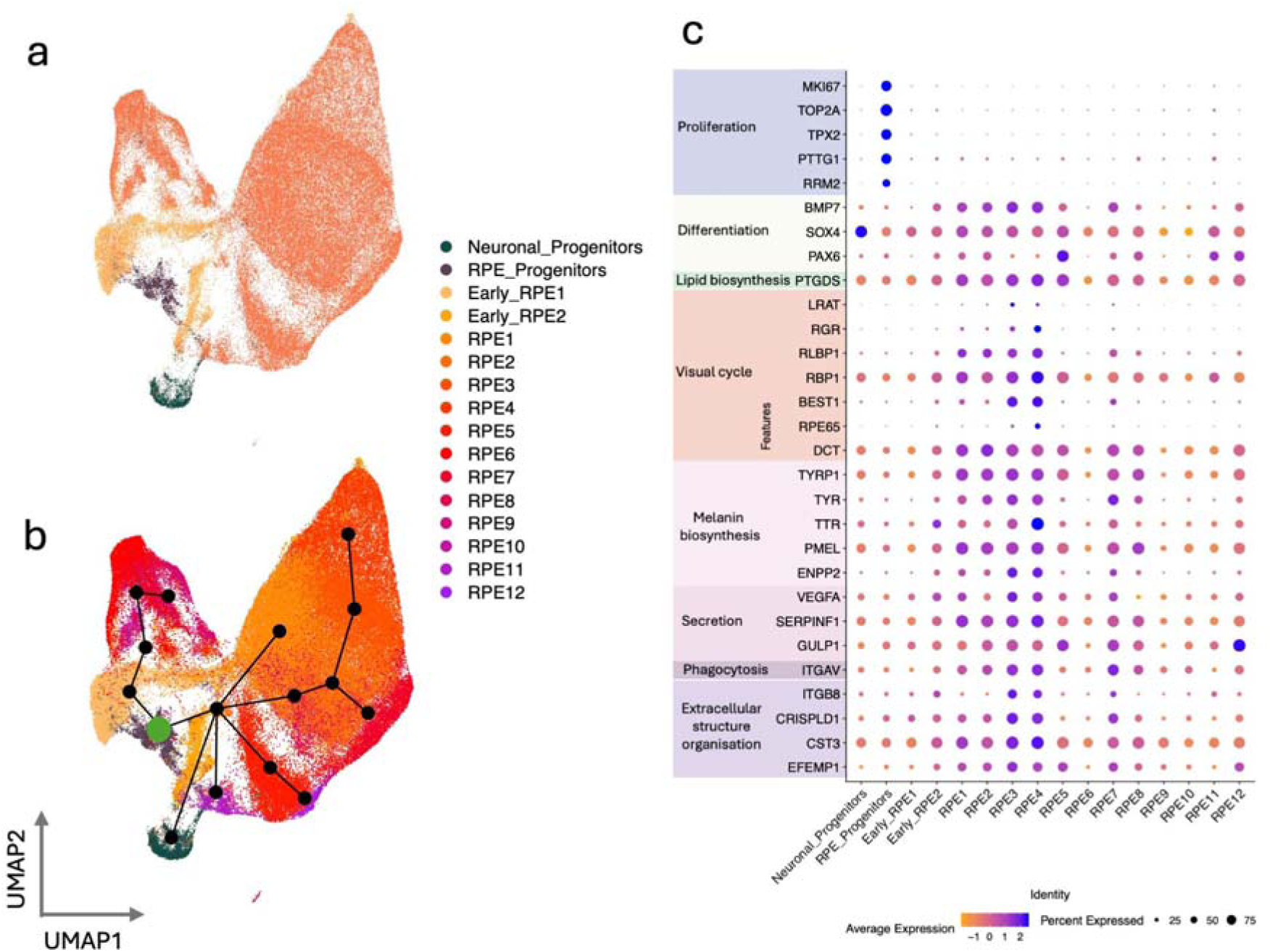
Characterisation of subpopulations. (**a**) Uniform Manifold Approximation and Projection (UMAP) of cells labeled by subpopulation identity and (**b**) showing cell trajectory. (**c**) Dotplot showing scaled average expression (z-scores; colour scale) and percentage of expressing cells (dot size) for genes associated with RPE functions (extracellular structure organization, phagocytosis, secretion, melanin biosynthesis, visual cycle, lipid biosynthesis, differentiation, and proliferation) across subpopulations.

**Table 1.** Number and % of cells per subphenotypes and sub-populations.

| Subpopulation | AMD/<br>RPD- | AMD/<br>RPD+ | Total | % AMD/<br>RPD- | % AMD/<br>RPD+ |
| --- | --- | --- | --- | --- | --- |
| Early RPE1 | 6,476 | 4,755 | 11,231 | 9.68 | 7.47 |
| Early RPE2 | 3,649 | 2,679 | 6,328 | 5.46 | 4.21 |
| Neuronal Progenitors | 1,549 | 1,273 | 2,822 | 2.32 | 2.00 |
| RPE Progenitors | 2,081 | 1,661 | 3,742 | 3.11 | 2.61 |
| RPE1 | 10,600 | 11,223 | 21,823 | 15.85 | 17.63 |
| RPE2 | 8,756 | 8,450 | 17,206 | 13.09 | 13.28 |
| RPE3 | 7,324 | 6,594 | 13,918 | 10.95 | 10.36 |
| RPE4 | 6,185 | 6,026 | 12,211 | 9.25 | 9.47 |
| RPE5 | 6,225 | 3,997 | 10,222 | 9.31 | 6.28 |
| RPE6 | 3,977 | 5,695 | 9,672 | 5.95 | 8.95 |
| RPE7 | 3,504 | 3,407 | 6,911 | 5.24 | 5.35 |
| RPE8 | 3,220 | 2,793 | 6,013 | 4.81 | 4.39 |
| RPE9 | 1,094 | 2,915 | 4,009 | 1.64 | 4.58 |
| RPE10 | 600 | 1,198 | 1,798 | 0.90 | 1.88 |
| RPE11 | 1,085 | 658 | 1,743 | 1.62 | 1.03 |
| RPE12 | 557 | 318 | 875 | 0.83 | 0.50 |
| Total | 66,882 | 63,642 | 130,524 | 100 | 100 |

### Differential gene expression between AMD/RPD+ and AMD/RPD− RPE

Differential gene expression analysis of single-cell RPE profiles using MAST (**Supplementary Data**) identified 728 genes that differed significantly between AMD/RPD+ and AMD/RPD− samples after multiple-testing correction (adjusted *p* < 0.05). Of these, 458 genes were upregulated and 270 downregulated in AMD/RPD+ relative to AMD/RPD− (**Figure 2**).

**Figure 2:**
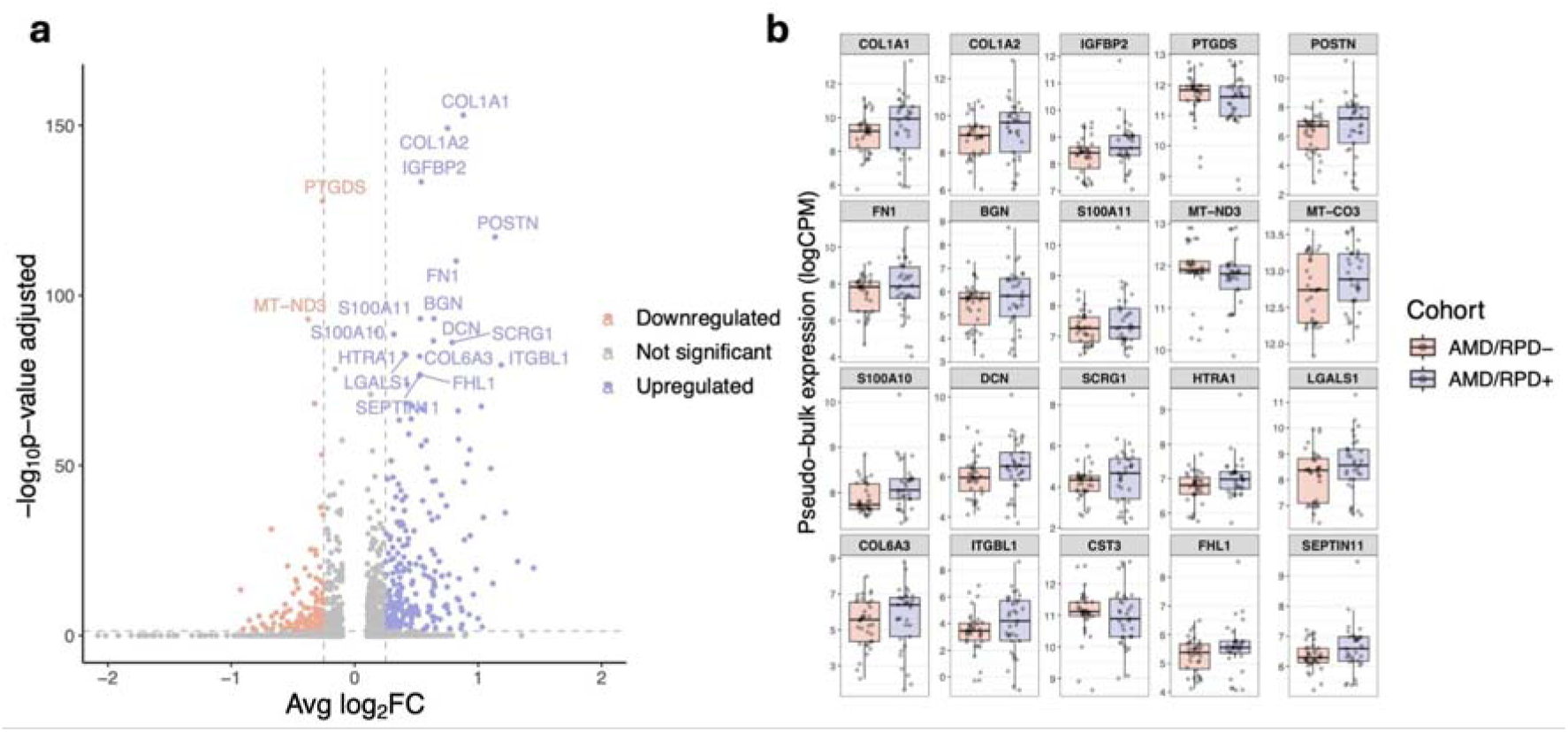
Differential gene expression in RPE cells from AMD/RPD+ and AMD/RPD− participants. (**a**) Volcano plot showing differentially expressed genes between RPE cells derived from AMD/RPD+ and AMD/RPD− participants (directionality relative to AMD/RPD+). Genes significantly downregulated in AMD/RPD+ (adjusted *p* < 0.05, log□FC > 0.25) are shown in red, and upregulated genes in blue. The top 20 differentially expressed genes are annotated. (**b**) Box plot of top 20 genes showing pseudobulk expression in logCPM (counts per million) across AMD/RPD+ and AMD/RPD− RPE samples.

Compared to the AMD/RPD− cohort, the AMD/RPD+ cohort showed upregulation of genes involved in ECM remodelling and structural integrity, including collagen (e.g. *COL1A1*, *COL1A2*), *FN1*, *POSTN*, *HAPLN1*, *ITGBL1,* and *HSPG2*. Upregulation of *LINC01133* and *VIM* suggests partial epithelial-mesenchymal transition and cytoskeletal reorganisation. The canonical AMD-associated gene *HTRA1* was also higher in AMD/RPD+, consistent with its role in ECM regulation and AMD progression [44,45]. Because variation at the *ARMS2/HTRA1* locus (rs11200638) is a major genetic risk factor for AMD, we compared genotype frequencies between groups: in the AMD/RPD− cohort, 45% of participants were heterozygous and 20% homozygous carriers, whereas in AMD/RPD+ 45% were heterozygous and 42% homozygous carriers (*p* = 0.04518). Thus, although genotype differences at *ARMS2/HTRA1* could contribute to the observed transcriptional differences, they are unlikely to fully explain them, as none of the significantly differentially expressed genes are known direct targets of this locus. Additional increases in mitochondrial (*MT-CYB*) and glycolytic genes (*LDHA*, *PGK1*, *FAM162A*), as well as *IGFBP2*, indicate metabolic adaptation to microenvironmental stress (**Supplementary Data**).

Conversely, genes upregulated in AMD/RPD− compared to AMD/RPD+ included multiple mitochondrial transcripts such as *MT-ATP8* (Complex V), associated with mitochondrial disorders, including retinitis pigmentosa [46], and NADH-dehydrogenase subunits *MT-ND2*, *MT-ND3*, *MT-ND4L*, *MT-ND5*, *MT-ND6*, which have been implicated in Leber Hereditary Optic Neuropathy [47], alongside nuclear-encoded Complex I components (*NDUFA1*, *NDUFA3, NDUFA4, NDUFA13*, *NDUFB1*, *NDUFB2*, *NDUFS7*). Stress-response genes (*HSP90B1*, *HSPE1*), both of which participate in protein folding and endoplasmic reticulum stress responses and implicated in AMD pathogenesis [48,49], as well as the antioxidant enzyme *GPX4* and *PTGDS* (prostaglandin D□ synthase) were also elevated, pointing to relative preservation or compensatory activation of mitochondrial and antioxidant pathways (**Supplementary Data**).

Over-representation analysis was used to assess pathway and disease associations of differentially expressed genes between the two cohorts [50], using the Gene Ontology [51] and Disease Ontology [52] databases (**Figure 3**). In AMD/RPD+, enriched biological processes involved ECM organisation, cytoskeletal remodelling, cell substrate adhesion, wound healing, and responses to hypoxia and oxygen levels (**Figure 3a**). Molecular function analysis showed enrichment for genes encoding proteins involved in the ECM, actin binding, integrin binding, collagen binding, and growth factor interactions, consistent with dynamic regulation of the extracellular environment and cellular attachment (**Figure 3b**). Enriched cellular components included the collagen-containing ECM, focal adhesion, and actin cytoskeletal structures, supporting a shift toward structural remodelling and mechanosensing (**Figure 3c**). KEGG pathways further highlighted over-representation of cytoskeleton, focal adhesion and ECM-receptor interaction pathways (**Figure 3d**). Disease Ontology enrichment revealed genes upregulated were associated with conditions involving connective tissue remodelling, vascular pathology, and cancer-related processes (**Figure 3e**).

**Figure 3:**
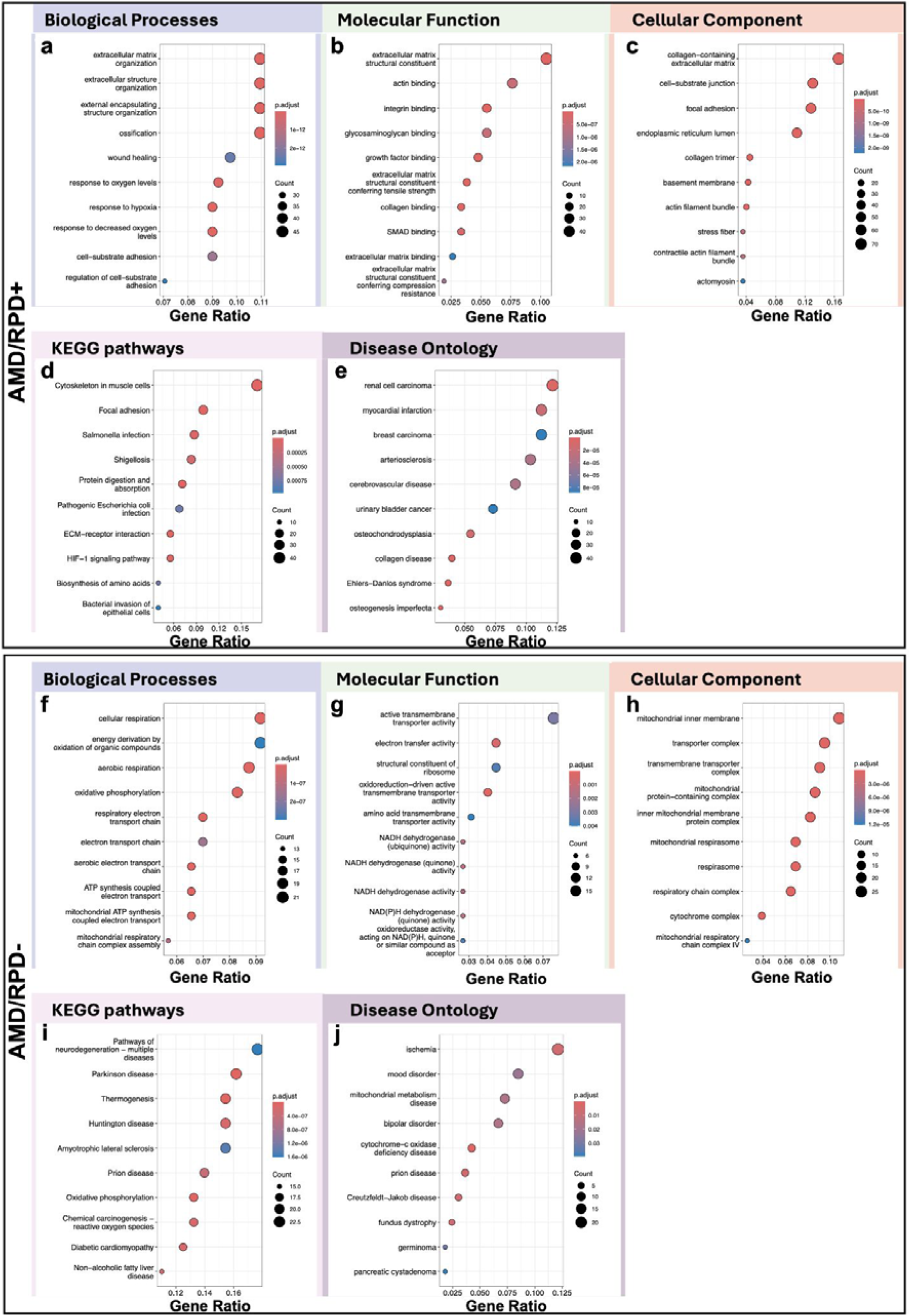
Enrichment analysis in each cohort. Pathway analysis showing the top 10 enriched Gene Ontology terms, KEGG pathways, and Disease Ontology categories for genes upregulated in RPE cells from (**a-e**) AMD/RPD+ compared to AMD/RPD−, and (**f-j**) AMD/RPD− compared to AMD/RPD+cohorts. Panels show the top enriched terms for (**a, f**) biological processes, (**b, g**) molecular functions, (**c, h**) cellular components, (**d, i**) KEGG pathways, and (**e, j**) disease ontology categories. Pathways are ranked by gene ratio and adjusted *p* value. Bubble size represents the number of genes associated with each term, and colour indicates the adjusted *p* value (p.adjust).

In AMD/RPD− RPE cells, enrichment was observed in mitochondrial energy metabolism, including cellular respiration, oxidative phosphorylation, electron transport chain activity, and ATP synthesis (**Figure 3f**). Genes involved in NADH-dehydrogenase, electron transfer and oxidoreduction activities were over represented, alongside terms for ribosome assembly (**Figure 3g**). Enriched cellular components included the mitochondrial inner membrane, the respirasome, the respiratory chain and cytochrome complexes (**Figure 3h**). KEGG pathways and Disease Ontology enrichment revealed associations with mitochondrial disorders (e.g. cytochrome c oxidase deficiency, oxidative phosphorylation), neurodegenerative diseases (e.g. Parkinson’s disease, Huntington’s disease, amyotrophic lateral sclerosis, Creutzfeldt-Jakob disease), and inherited retinal dystrophies (e.g. fundus dystrophy) (**Figure 3i, j**).

Collectively, these data define two distinct transcriptional states within AMD. AMD/RPD+ RPE cells engage ECM remodelling and hypoxia-adaptive programs. In contrast, AMD/RPD− cells display relative enrichment of mitochondrial, translational, and protein homeostasis processes, including oxidative phosphorylation and other energy-related pathways.

### Subpopulation-specific regulatory signatures in the RPE

To explore the impact of common genetic variation on gene regulation in the RPE, we conducted expression quantitative trait locus (eQTL) mapping within each subpopulation and identified 436 significant eQTLs (false discovery rate < 0.05). The number and distribution of eQTLs varied across cell subpopulations, with the highest burden observed in RPE3, RPE12, and RPE8 (**Table 2, Supplementary Data, Figures 4, S3**). We identified several eQTLs mapping to genes involved in diverse cellular processes, including signalling (*MAP2K2*), tRNA modification (*ELP5*), protein sorting (*RER1*), GPI-anchor remodeling (*PGAP2*), and lipid metabolism with links to oxidative stress (*TLCD5*). These eQTLs were not significantly associated with disease status. Because all individuals in both cohorts already had AMD, these eQTLs are unlikely to represent primary risk factors for disease onset. Instead, they could reflect genetic variation that contributes to phenotypic heterogeneity among patients. A subset of eQTLs showed significant interaction with disease status (SNP:disease *p* < 0.05), implicating loci with regulatory effects that may differ in AMD (**Table 2**, **Figures 4, S3**). These include *AP3S1* (RPE3), which encodes an adaptin protein of the AP-3 complex involved in intracellular trafficking, including cargo sorting from the Golgi apparatus to lysosomes [53], as well as melanocyte trafficking and melanogenesis [54]. This is of particular interest in the RPE, where such pathways are essential for melanosome biogenesis and lysosomal clearance of photoreceptor outer segments, processes linked to drusen formation and AMD pathology [55]. Other notable eQTLs include *AIPL1* (RPE1), already associated with inherited retinopathies [56]; *FSTL5* (RPE8), a follistatin-like gene with enriched retinal expression, implicated in BMP signaling, ECM interactions, and development; and the developmental transcription factors *MSX2* (RPE3) and *ZFPM2* (RPE4). Signals in developmental regulators may reflect stressed RPE cells re-engaging differentiation programs, although some activity could also arise from residual immaturity of iPSC-derived RPE cells. *MC5R*, a G-protein-coupled receptor involved in melanocortin signalling and immune modulation, showed disease-associated regulation in neuronal progenitors, pointing to possible neuroimmune mechanisms influencing AMD susceptibility. Additional disease-interacting eQTLs mapped to long non-coding RNAs, suggesting roles for post-transcriptional or epigenetic regulation (**Table 2**). Finally, we investigated whether previously reported SNPs influencing *ARMS2/HTRA1* expression, including variants recently associated with RPD (rs11200638, rs79641866/PARD3B, rs143184903/ITPR1, rs76377757/SLN, and the lncRNA gene *HTRA1-AS1*) were associated with altered gene expression in our dataset. rs11200638 was directly tested against both *HTRA1* and *ARMS2* but was not significant. The other reported SNPs were not present in our reference panel or were filtered during imputation quality control. However, nearby variants were tested (rs849124 near *PARD3B*, rs4234561 near *ITPR1*, and rs4754244 near *SLN*) and none were significant. No variants were significantly associated with *HTRA1-AS1* expression. These results indicate that known AMD-associated and RPD−associated SNPs do not explain the elevated HTRA1 transcript and protein levels observed in the AMD/RPD+ RPE cohort. In summary, eQTLs capture both baseline regulatory variation and disease-specific effects, underscoring genetic contributions to differences in AMD subphenotypes.

**Figure 4.**
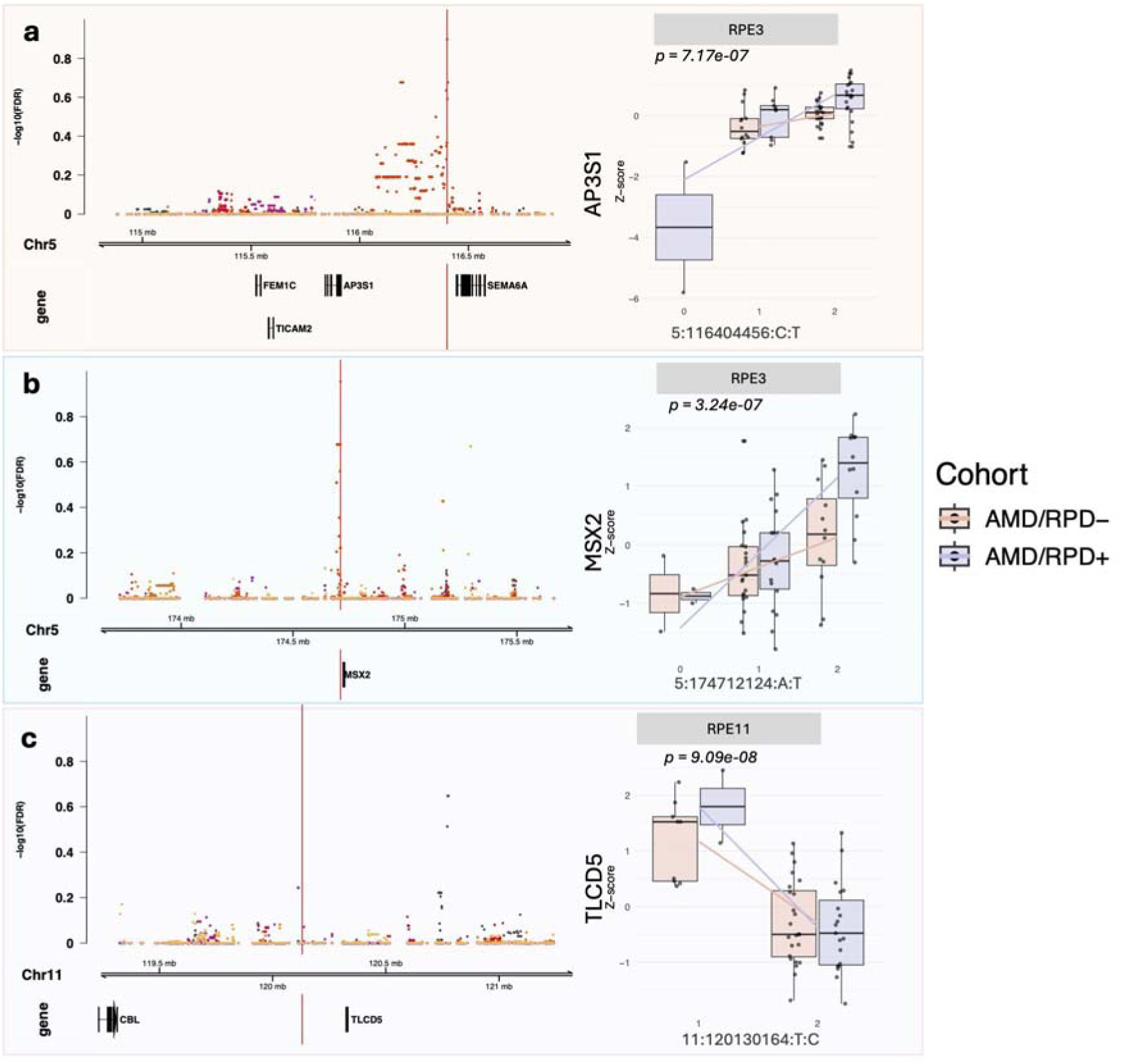
Examples of disease-interacting eQTLs in RPE subpopulations. Regional association plots (left) depict local association signals (-log10 false discovery rate) across the genomic locus, with the lead SNP indicated by a vertical red line. Boxplots (right) show expression stratified by genotype and cohort, with regression lines illustrating allele dosage effects. (**a**) *AP3S1* in RPE3 (5:116404456 C>T; *p* = 7.17 × 10□□). (**b**) *MSX2* in RPE3 (5:174712124 A>T; *p* = 3.24 × 10□□). (**c**) *TLCD5* in RPE11 (11:120130164 T>C; *p* = 9.09 × 10□□).

**Table 2.**
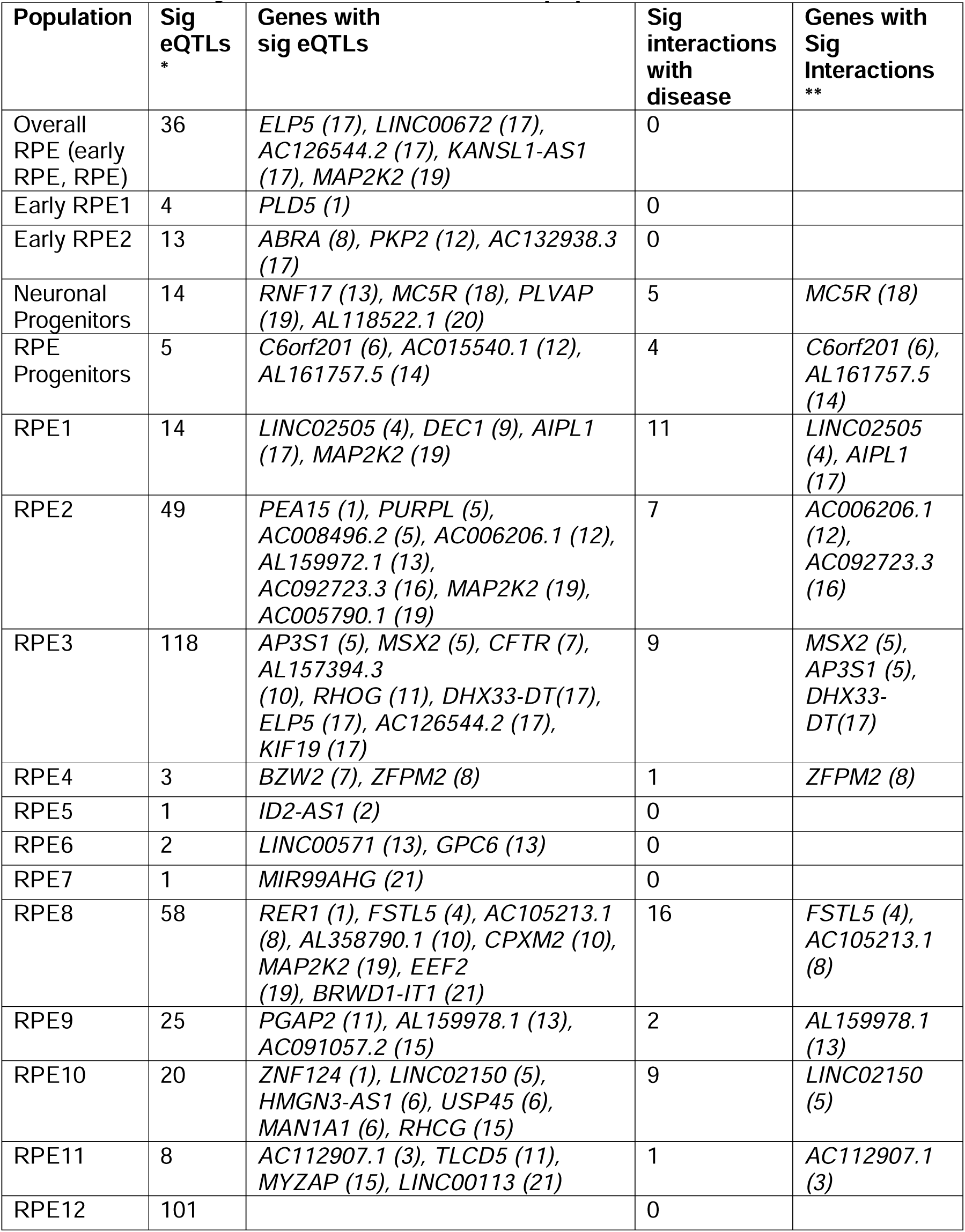
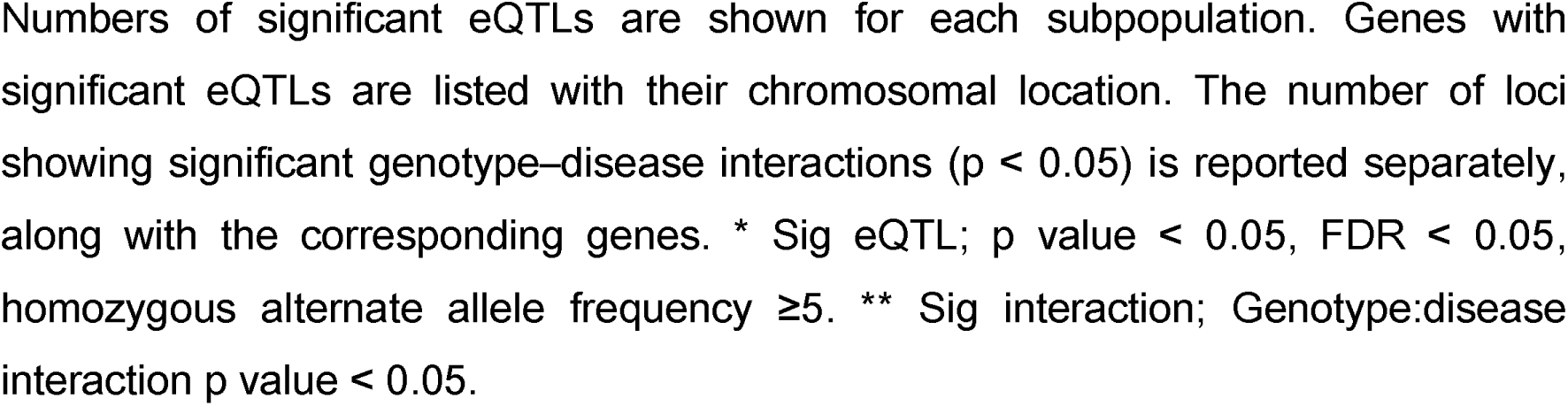
Summary of eQTLs across RPE subpopulations.

### Differential protein expression between AMD/RPD+ and AMD/RPD− RPE

To complement the transcriptomic analysis, we profiled the proteome of RPE cultures using tandem mass tag (TMT)-based mass spectrometry on bulk protein lysates. A total of 5,849 proteins were identified across six TMT runs, filtered at a 1% false discovery rate (**Supplementary data**). Differentially expressed proteins were identified using standard statistical thresholds, and functional enrichment was performed using STRING v12.0 [57]. In total, 715 proteins were significantly differentially abundant between cohorts, with 502 upregulated and 213 downregulated in AMD/RPD+ relative to AMD/RPD− (**Figures 5a, 5b, S4**). Of these, 35 upregulated and 27 downregulated proteins have previously been associated with RPE dysfunction or AMD pathology. Canonical RPE markers were similarly expressed between groups (**Figure S1)**. The top 16 differentially abundant proteins are shown in **Figure S4** for illustration, highlighting representative up- and down-regulated species across biological replicates.

**Figure 5.**
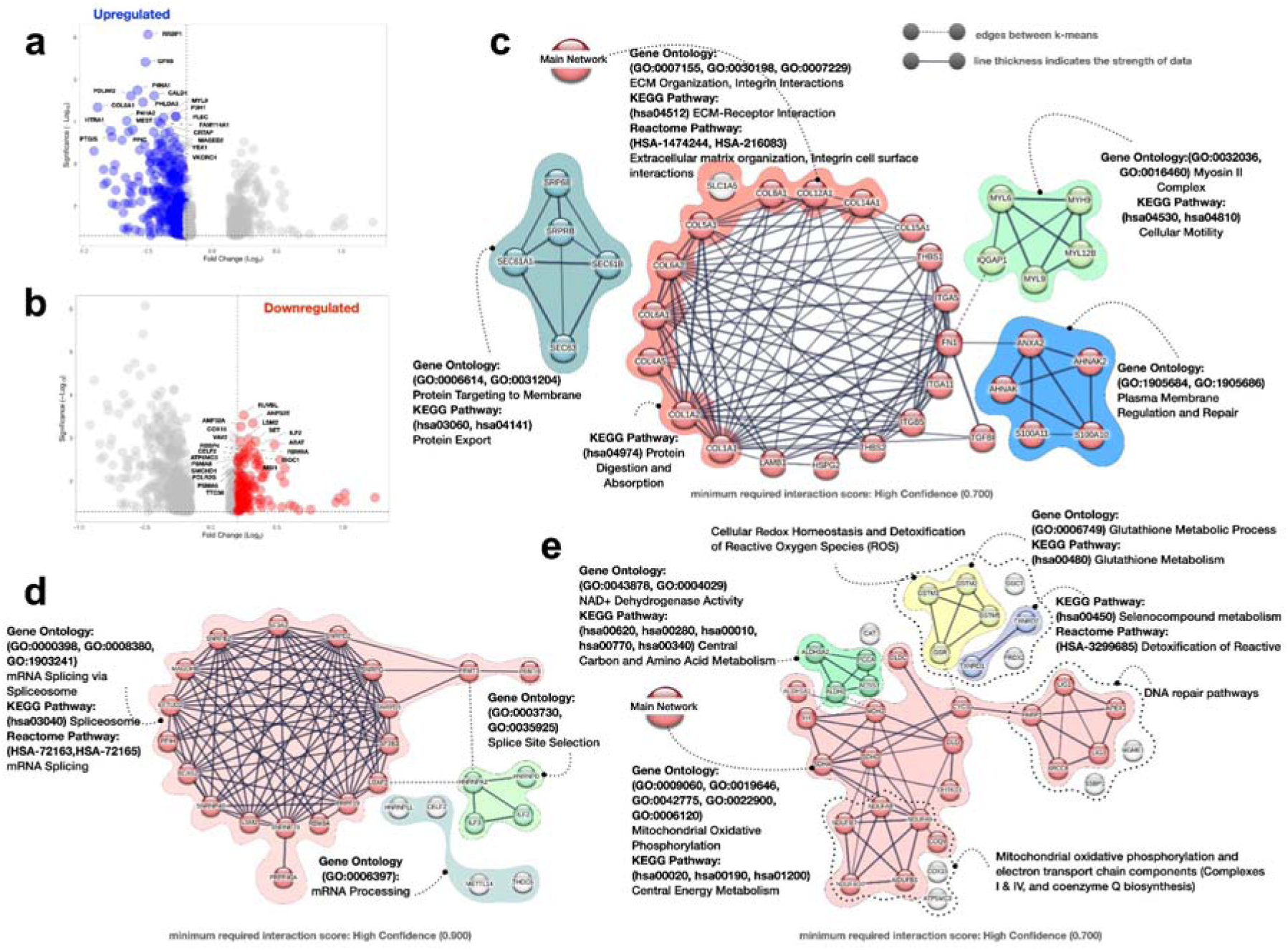
Differential protein expression between AMD/RPD+ and AMD/RPD−RPE cells. (**a, b**) Volcano plots showing proteins significantly upregulated (a, blue) and downregulated (b, red) in AMD/RPD+ relative to AMD/RPD− RPE cells (false discovery rate < 0.05, log□ fold-change > 0.5). Selected proteins of interest are annotated. (**c-e**) Protein-protein interaction networks of differentially abundant proteins identified by STRING (v12.0). (**c**) Upregulated proteins in AMD/RPD+ form enriched clusters related to extracellular matrix (ECM) organisation, cytoskeletal regulation, and membrane signalling. (**d-e**) Downregulated proteins in AMD/RPD+ (upregulated in AMD/RPD−) cluster in modules related to mitochondrial metabolism, oxidative phosphorylation, RNA splicing, and antioxidant defence. (**a-e**) Node size represents protein abundance; edge thickness corresponds to interaction confidence (minimum interaction score > 0.7). Functional clusters are colour-coded according to pathway enrichment (see **Supplementary Data)**.

Many proteins were significantly upregulated in the AMD/RPD+ cohort. These formed coherent clusters involved in ECM organisation, cytoskeletal regulation, and membrane signalling (**Figure 5c**). Notable ECM-associated proteins included COL1A1, COL1A2, COL4A5, COL6A2, FN1, and HSPG2, many of which are critical for matrix integrity and cell-ECM interactions. KEGG pathway analysis highlighted ECM-receptor interaction, protein digestion and absorption, and protein export, involving proteins such as SEC61A1, SEC61B, and SRP68. Increased expression extended to receptors for collagen and fibronectin, including ITGA11, associated with the blood-retinal barrier [58] and ITGB5, necessary for binding photoreceptor outer segments [59]. Components of the myosin motor complex (e.g. MYH9, MYL6, MYL9) were also increased, suggesting changes in cellular motility and structural tension. Among the 20 most significantly upregulated proteins in the AMD/RPD+ group were several previously linked to AMD pathogenesis (VKORC1 [60], PLEC [61], COL8A1 [62,63], HTRA1 [45]) or retinal disease (CALD1 [64], MYL9 [65], P3H1 [66], P4HA2 [67], RRBP1 [68]). Proteins involved in membrane integrity and calcium signalling, such as ANXA2, one of the most common proteins found in drusen and linked to inflammatory disease states [69,70], as well as S100A10 and AHNAK, both associated with membrane repair, were also increased in abundance.

Several proteins were significantly reduced in the AMD/RPD+ cohort compared with AMD/RPD− (more abundant in the AMD/RPD− cohort, **Figure 5a, Supplementary data**). Among those were proteins previously linked to glaucoma (ANP32A [71], VAV2 [72]) and retinal biology (MSI1 [73], RBM8A [74]). Functional enrichment revealed increased representation of pathways related to RNA splicing and mRNA processing, reflected by elevated abundance of spliceosome-associated proteins (e.g. SNRNP70, RBM8A, PRPF19, U2AF2, SF3B3, HNRNPA1, CELF2) (**Figure 5d)**. Some of these have been implicated in retinal degenerative conditions (e.g. SNRPD1, SNRPD2, SNRPG, SNRPB2, SF3A2) [72,73] Pathways linked to DNA repair and cellular maintenance were also less represented in the AMD/RPD+ cohort (**Figure 5e**), including proteins involved in base excision repair (e.g. PARP1, LIG1, LIG3, APEX1) and antioxidant defence (e.g. PRDX2, GSR, CAT). Multiple aldehyde dehydrogenases (e.g. ALDH2, ALDH3A2, ALDH1A1) showed lower abundance, suggesting reduced capacity to detoxify reactive aldehydes and to mitigate lipid peroxidation (**Figure 5e**). Similarly, mitochondrial metabolism pathways were downregulated in AMD/RPD+, with decreased abundance of TCA cycle enzymes (DLD, MDH2, FH) and glutathione metabolism proteins (GSTM2, GSTM3, GSTM5, GGCT), indicating diminished antioxidant and bioenergetic capacity compared to the AMD/RPD− cells (**Figure 5e**). Consistent with the transcriptomic findings, proteins involved in mitochondrial respiration and oxidative phosphorylation were also reduced in AMD/RPD+ (**Figure 5e**). These include Complex I subunits (NDUFA8, NDUFA9, NDUFB3, NDUFB10), cytochrome oxidase biosynthesis (COX15), and electron transport components (COQ9, ATP5MC3). Several of these proteins have previously been implicated in the pathophysiology of geographic atrophy in RPE cells [41].

### Protein-level regulatory signatures in the RPE

To explore the genetic regulation of protein abundance, we performed protein quantitative trait locus (pQTL) analyses. We identified five significant pQTLs present within both AMD/RPD− and AMD/RPD+ cohorts (false discovery rate < 0.05; **Tables 3, S1, Figure 6**). No differences were observed between AMD/RPD− and AMD/RPD+ cohorts. Four pQTLs mapped to proteins with mitochondrial functions, including KYAT3 (tryptophan metabolism, producing kynurenic acid with neuroprotective roles, including in the retina [75,76]), PYROXD2 (oxidoreductase, already associated with geographic atrophy in RPE cells [41]), GLRX5 (iron-sulfur cluster biogenesis, with impaired metabolism reported in a retinal organoid model of autosomal dominant retinitis pigmentosa under ATF6 stress [77]), and NQO1 (redox cycling, also already implicated in AMD [78]), underscoring a consistent role for mitochondrial pathways across molecular layers. The only non-mitochondrial hit was AP4B1, an adaptor protein complex subunit involved in intracellular trafficking. Notably, AP4B1 mutations cause autosomal recessive spastic paraplegia, a neurodegenerative disorder that can present with visual dysfunction [79], implicating lysosomal trafficking in retinal health. Both GLRX5 and AP4B1 converge on cellular iron regulation, GLRX5 via mitochondrial Fe-S cluster biogenesis and AP4B1 via endo-lysosomal trafficking pathways that influence iron handling. This suggests iron homeostasis is a potential unifying axis in AMD pathobiology. Together, these findings indicate that mitochondrial regulation forms a shared genetic backbone of AMD. At the same time, the distinction between RPD−positive and RPD−negative disease likely reflects additional regulatory mechanisms beyond baseline protein-level genetic effects.

**Figure 6.**
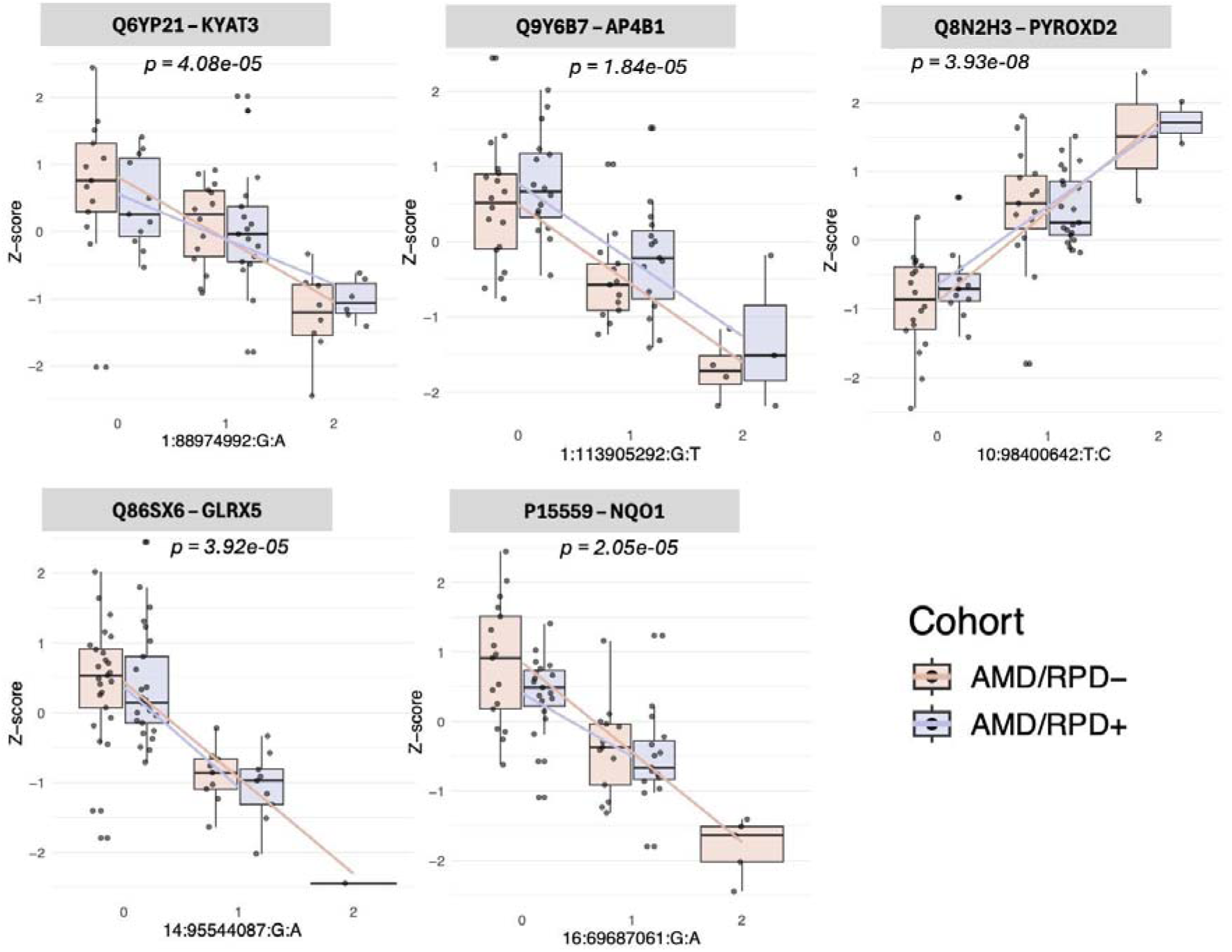
Protein quantitative trait loci in RPE cells. Representative pQTLs showing association between genotype and protein abundance for KYAT3 (Q6YP21), AP4B1 (Q9Y6B7), PYROXD2 (Q8N2H3), GLRX5 (Q86SX6), and NQO1 (P15559). Boxplots display rank-normalized protein abundance (y-axis) stratified by genotype (x-axis), with data shown separately for AMD/RPD− (red) and AMD/RPD+ (blue) cohorts. Linear regression fits are overlaid to illustrate allele dosage effects.

**Table 3.**
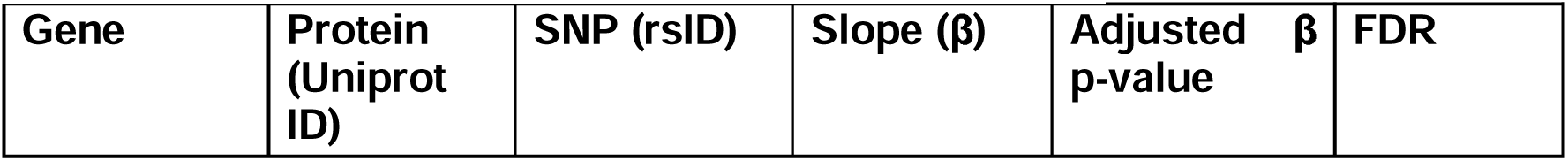
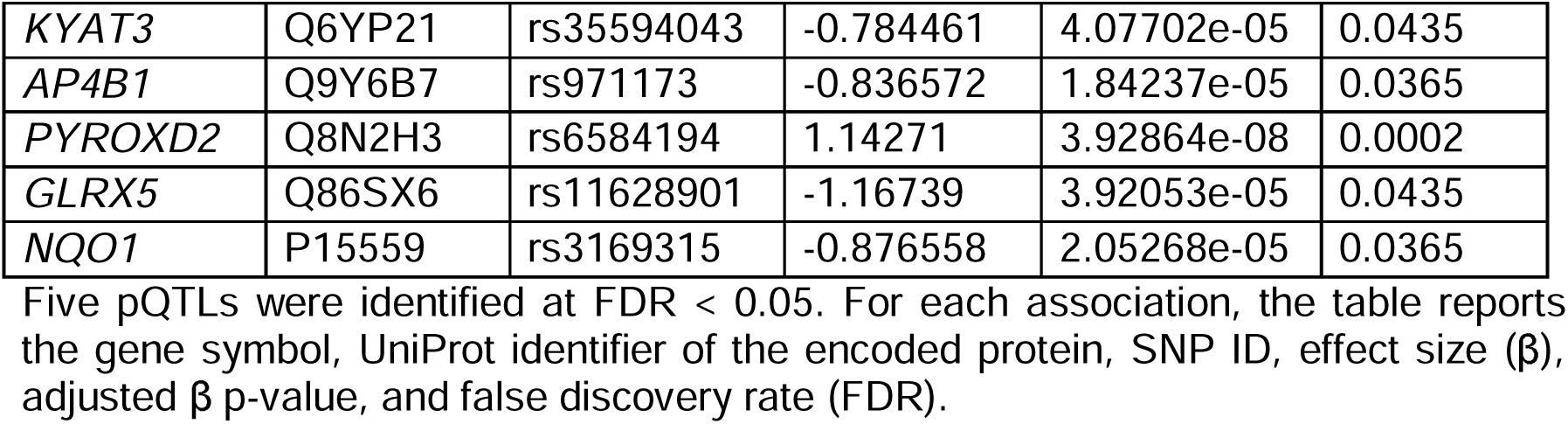
Significant protein quantitative trait loci.

### Integrating genetic regulation with disease risk

To further integrate genetic and transcriptomic signals, a transcriptome-wide association study (TWAS) was performed as a complementary approach to eQTL and pQTL analyses, linking genetic variation to disease through predicted gene expression and thereby prioritising candidate genes beyond individual QTL effects. Sixty gene-trait associations reached significance across the overall RPE population and subpopulations (**Figure 7, Supplementary data**). Genes related to mitochondrial metabolism and homeostasis (*LYRM4*, *ALDH9A1*, *EGLN3/PHD3*, *GPS2*) were associated with RPD risk. *LYRM4* encodes ISD11, an iron-sulfur cluster biogenesis factor whose loss-of-function mutations cause combined mitochondrial respiratory chain deficiencies [80]; *ALDH9A1* contributes to aldehyde metabolism and redox balance; EGLN3/PHD3, a hypoxia-inducible prolyl hydroxylase regulating mitochondrial-dependent apoptosis, showed lower genetically predicted expression associated with lower RPD risk, consistent with a potential contribution of stress-induced cell-death pathways; and *GPS2*, a transcriptional coregulator that translocates from mitochondria to nucleus during stress to promote mitochondrial gene expression and maintain organelle homeostasis [81] (UniProt Q13227), suggests altered mitochondrial-nuclear communication may modulate susceptibility. In parallel, *SMAD3* and *TCF21*, which act within a shared TGF-β responsive network regulating mesenchymal differentiation and ECM transcription [82], were also associated with RPD. Notably, SMAD3 is required for RPE epithelial-mesenchymal transition and subretinal fibrosis following retinal detachment [83], underscoring its established involvement in retinal fibrotic responses. Among other significant loci, several map to genes with prior retinal relevance including *DGCR8* (RPE microRNA biogenesis; RPE survival and photoreceptor maintenance [84,85], *FEZF1* (postmitotic regulator of ON starburst amacrine cell fate [86]), *PSMC1* (26S proteasome ATPase supporting RPE–photoreceptor phagocytosis [87], and *TENM4* (axon guidance and retinal circuit development) [88]. These associations, interpreted with caution given TWAS limitations, point to inherited modulation of mitochondrial and ECM regulation in the RPE.

**Figure 7.**
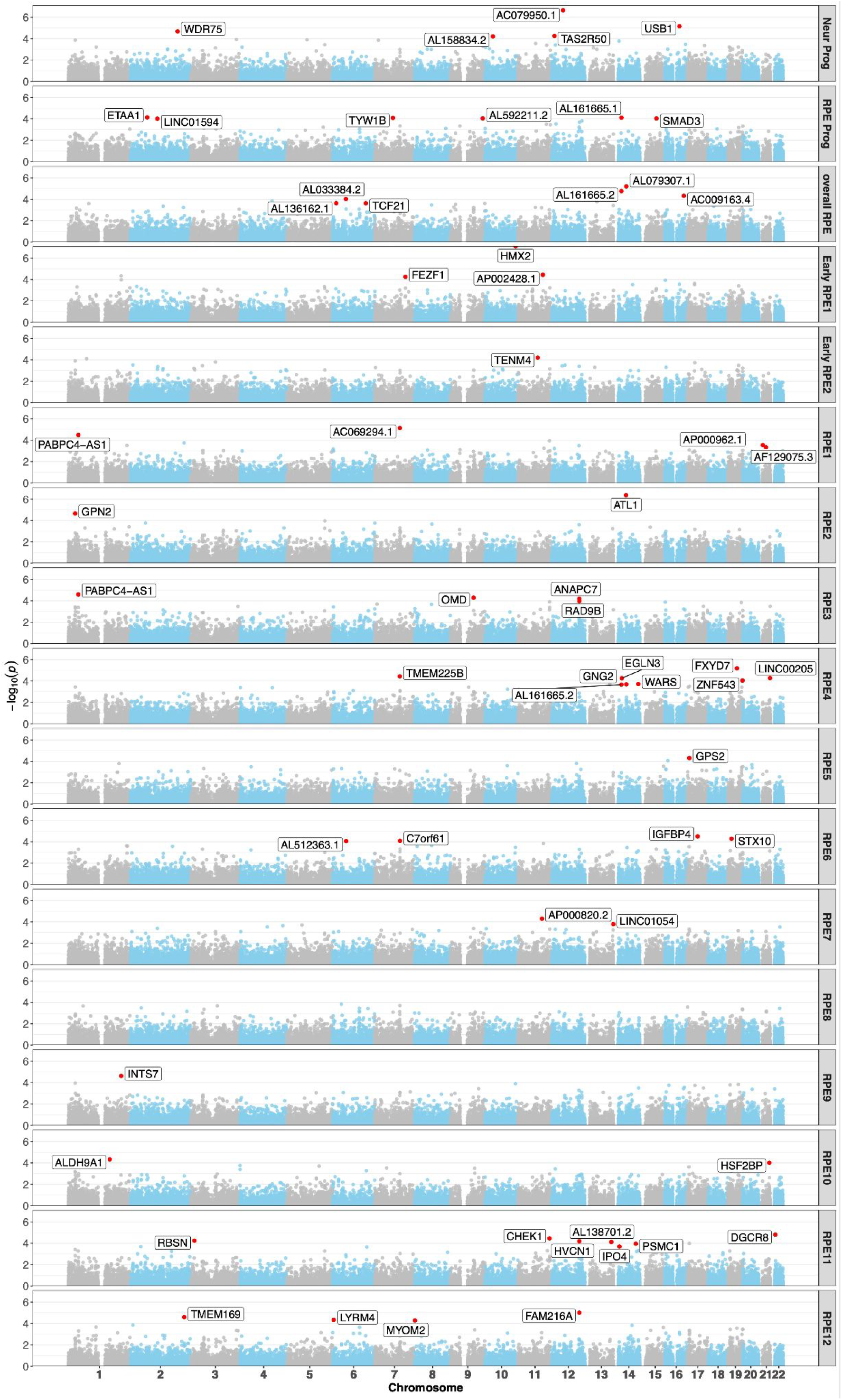
Transcriptome-wide association study (TWAS) of RPD risk across RPE populations. Manhattan plots showing TWAS results for overall and subpopulation-specific RPE transcriptomic models. Each point represents a gene-level association between genetically predicted expression and RPD risk, plotted by genomic position. The red dashed line denotes the significance threshold (false discovery rate < 0.1). Significant loci are labelled.

### Functional analyses

Both transcriptomic and proteomic analyses revealed differences in ECM organisation and cell-substrate adhesion pathways between AMD cohorts with and without RPD. However, transmission electron microscopy did not show consistent ultrastructural differences between the two groups (**Figure S5**). To test whether these molecular signatures corresponded to functional differences, we performed assays in iPSC-derived RPE cells generated from control donors (selected from our previous studies [41]), and from AMD/RPD− and AMD/RPD+ donors.

As we previously reported [89], all three iPSC-derived RPE cohorts formed drusen-like deposits *in vitro*. Among the AMD-derived cells, the AMD/RPD− lines produced more basal deposits than either the control or AMD/RPD+ groups (**Figure 8a-c, Supplementary Video**). To assess stress susceptibility, RPE cells were exposed to the toxic N-Retinylidene-N-Retinylethanolamine (A2E, 10 µM, 7 days), which accumulates with aging and AMD, and induces oxidative and lysosomal stress (**Figure 8d**). After treatment, AMD-derived RPE wells showed widespread detachment (absent in controls), most markedly in the AMD/RPD+ group (**Figure 8e**). Despite this structural instability, overt cell death was only observed in A2E-treated AMD/RPD− cultures, while AMD/RPD+ cells largely remained viable (**Figure 8f**). Because extensive detachment in AMD/RPD+ wells precluded accurate quantification, drusen-like deposits could not be measured in this group. However, A2E exposure significantly increased drusen-like deposits in the remaining attached AMD/RPD− cultures compared with controls (**Figure 8g, h**). Collectively, these findings suggest that the two AMD subphenotypes exhibit distinct modes of functional impairment: AMD/RPD+ cells display structural fragility and detachment under stress, whereas AMD/RPD− cells show greater metabolic vulnerability and cell loss. These differences mirror their respective molecular programs, with ECM remodelling dominating in one and mitochondrial and oxidative stress pathways in the other.

**Figure 8:**
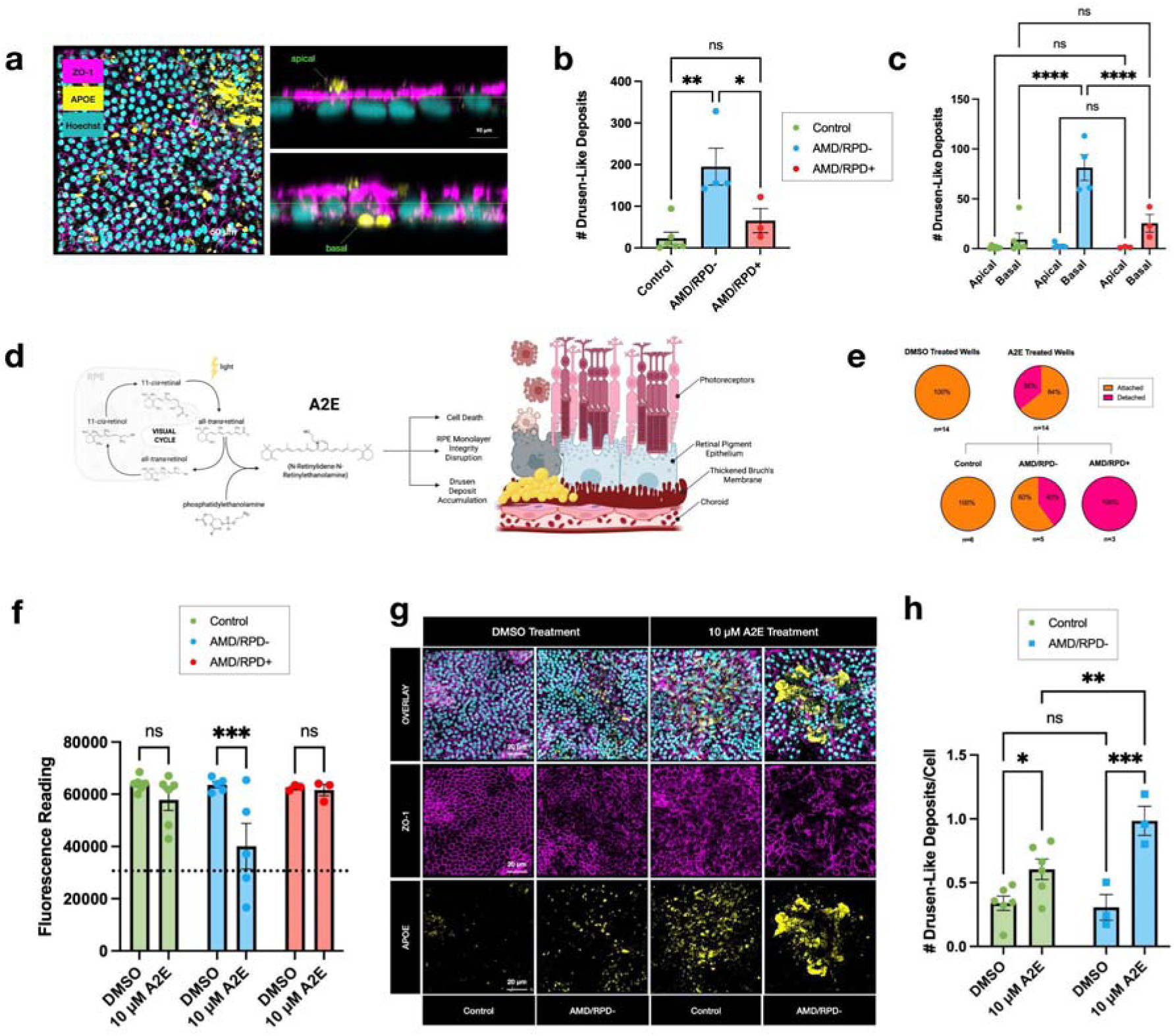
Differential drusen-like deposit formation between cohorts. (**a**) Left: representative high-resolution Z-stack ZO-1, APOE, Hoechst. Scale bar: 50 µm. Right: orthogonal view of Z-stack, lime horizontal bar to show apical versus basal distinction. Scale bar: 10 µm. (**b**) Number of drusen-like deposits in control and AMD lines. Data are mean ± SEM of triplicate values per well from all lines. Control: 9.1 ± 10.2, n=6 lines; AMD/RPD−: 195.2 ± 88.8, n=4 lines; AMD/RPD+: 65.4 ± 50.3, n=3 lines; p-values: AMD/RPD− versus AMD/RPD+: 0.0408, AMD/RPD− versus Control: 0.0030, AMD/RPD+ versus Control: 0.5897. (**c**) Number of apical versus basal drusen-like deposits. Mean deposit number <50 µm^3^ ± SEM: AMD/RPD−: apical: 3.0 ± 2.9, basal: 81.3 ± 25.9, n=4 lines; AMD/RPD+: apical: 1.2 ± 0.7, basal:25.4 ± 15.4, n=3 lines; Control: apical: 1.8 ± 1.6, basal: 2.7 ± 2.9 n=6 lines; p-values for basal drusen-like deposit comparison: AMD/RPD− vs. AMD/RPD+: <0.0001, AMD/RPD−vs. Control: <0.0001; AMD/RPD+ vs Control: 0.2390. (**d**) Schematic of the impact of A2E on RPE cells; build-up of A2E in the cells induces cell death, RPE monolayer integrity disruption, and drusen deposit accumulation. Created in BioRender. Hall, J. (2026) under CC BY 4.0 license agreement #: SW28ZE3T50 https://BioRender.com/cln2ln4. (**e**) Pie chart breaking down detachment in treated wells. (**f**) PrestoBlue viability readings. Mean fluorescence value ± SEM: AMD/RPD−: DMSO: 63592.0 ± 2560, A2E: 40113.8 ± 19513.8, n=5 lines; AMD/RPD+: DMSO: 62574.3 ± 1067.9, A2E: 61580.0 ± 3810.4, n=3; lines Control: DMSO: 63950.0 ± 2784.1, A2E: 57841.5 ± 9841.6, n=6 lines; p-values for DMSO versus A2E treatment: AMD/RPD−: <0.0010, AMD/RPD+: 0.9020, Control: 0.2910. Dotted line marks viability cutoff; samples below this line were excluded in drusen-like deposit analysis in (h). (**g**) DMSO- and A2E-treated RPE exhibiting tight junction marker ZO-1 (pink), drusen deposit marker APOE (yellow) and nuclear stain Hoechst. Scale bar: 20 µm. (**h**) Number of drusen-like deposits normalized to cell count. Mean normalized deposit number ± SEM: AMD/RPD−: DMSO: 0.3 ± 0.2, A2E: 1.0 ± 0.2, n=3; Control: DMSO:0.3 ± 0.1, A2E: 0.6 ± 0.3 n=6 lines; p-values for drusen-like deposit comparison: Control: DMSO vs. A2E: 0.0337; AMD/RPD−: DMSO versus A2E: 0.0008; Control vs. AMD/RPD−: DMSO: 0.9449, A2E: 0.0080. (**b, c, f, h**) Statistical significance was established by 2-way-ANOVA followed by Tukey’s Multiple Comparisons Test, **** *p*<0.0001; *** *p*<0.001; ** *p*<0.01; * *p*<0.05.

## Discussion

This study delineates molecular and functional differences between two clinically defined subphenotypes of AMD: AMD/RPD− and AMD/RPD+. All differential expression and QTL analyses included age and sex as covariates to mitigate confounding by these factors. AMD/RPD+ RPE cells display transcriptional and proteomic signatures of ECM remodelling, cytoskeletal reorganisation, and microenvironmental adaptation, reflected by enrichment of hypoxia-responsive and adhesion pathways, together with relative depletion of mitochondrial and protein homeostasis pathways compared to AMD/RPD− cells. Importantly, both subphenotypes show molecular evidence of mitochondrial dysfunction and disrupted protein homeostasis, consistent with AMD being characterised by impaired mitochondrial regulation, but they differ in the relative prominence of these pathways. This suggests that RPD represents a distinct pathological trajectory that is more influenced by extracellular or microenvironmental dysregulation.

Despite sharing features such as drusen, AMD patients with RPD are clinically recognised to have worse visual function and a greater risk of progression to late-stage AMD than patients without RPD [10,14]. RPD has been associated with alterations in macular structure and retinal integrity, which have been linked to functional impairment in several studies [10]. Such tissue-level alterations could modify signalling and metabolic interactions between the neurosensory retina and the RPE, creating conditions that favour hypoxia-responsive and ECM-remodelling programs. This is consistent with our observation that AMD/RPD+ RPE cells activate structural, hypoxia-responsive and ECM-related pathways. This also likely reflects intrinsic differences in AMD/RPD+ RPE cells that persist despite reprogramming and differentiation in a shared *in vitro* environment, supporting the idea that donor-specific transcriptional wiring is retained. It may also help explain why, under our *in vitro* conditions, AMD/RPD+ RPE cells do not show overt metabolic failure despite molecular signatures of mitochondrial dysfunction and disrupted protein homeostasis, and suggests that disease drivers in RPD may, at least in part, reside outside the RPE, in the tissue context absent from monocultures.

Integration of genetic data further sharpens this distinction. Across cohorts and modalities, including eQTL, pQTL and TWAS analyses, we observed shared mitochondrial regulation in both subphenotypes, with comparatively greater mitochondrial vulnerability in AMD/RPD− and enrichment of ECM and structural programs in AMD/RPD+. While absolute expression levels cannot be inferred without direct comparison to controls, our previous work in geographic atrophy RPE demonstrated mitochondrial impairment relative to healthy RPE [41], consistent with metabolic vulnerability across subphenotypes. We identified convergent genetic regulation of mitochondrial and iron-related pathways (e.g. GLRX5, AP4B1, KYAT3, NQO1), consistent with these processes contributing to shared aspects of AMD susceptibility. In contrast, disease-interacting eQTLs highlighted subtype-specific regulation in RPD, including loci involved in intracellular trafficking, developmental transcription factors, and long non-coding RNAs. The additional association with proteasome-related genes is consistent with a possible involvement of phagocytic clearance mechanisms at the RPE-photoreceptor interface, and raises the possibility that altered protein homeostasis could contribute to inefficient debris removal in RPD.

Taken together, these multi-omics findings support a layered model in which baseline genetic risk establishes mitochondrial and iron-related vulnerability common to AMD. Additional regulatory mechanisms, particularly in ECM, cytoskeleton and adhesion pathways, and in aspects of protein homeostasis, distinguish AMD with RPD from conventional AMD. These partly separable trajectories may result in atrophy development through different initiating mechanisms. This aligns with existing models of AMD pathogenesis that emphasise RPE dysfunction as a primary initiator of disease in conventional AMD [90] and is compatible with the hypothesis that AMD/RPD+ is more strongly shaped by extrinsic tissue context.

## Conclusions

These differences have practical implications. First, our findings challenge the view that the presence of RPD represents a cumulative extension of conventional AMD pathology. Second, it highlights the limitations of AMD classifications based solely on conventional drusen. Integrating molecular profiling into AMD subclassification may improve prognostic resolution and therapeutic targeting. Finally, the findings underscore the strengths and caveats of iPSC-derived RPE cell modelling. While these effectively capture cell-intrinsic responses, they do not reflect the influence of surrounding retinal, vascular, or immune components that are likely critical in RPD formation.

In summary, this study defines two distinct molecular phenotypes within AMD at the level of iPSC-derived RPE. These findings advance the understanding of AMD heterogeneity and support a shift toward mechanism-based classification, with potential implications for future therapeutic stratification tailored to the distinct trajectories of AMD with and without RPD.

## Methods

### Participant recruitment

All participants were over 50 years old and gave informed written consent. This study was approved by the Human Research Ethics committees of the Royal Victorian Eye and Ear Hospital (20/1459H; 11/1031H) as per the requirements of the National Health and Medical Research Council (NHMRC), in accordance with the Declarations of Helsinki and with the International Conference on Harmonisation guidelines for Good Clinical Practice. Cases were recruited through ongoing AMD natural history studies at the Centre for Eye Research Australia as follows: 1) AMD with only conventional drusen (AMD/RPD−, 56 individuals, 42 females, mean ± SD age at recruitment: 70.8 ± 7.9 years); and 2) AMD cases with both conventional drusen and with extensive RPD (AMD/RPD+, 57 individuals, 35 females, mean ± SD age at recruitment 78.6 ± 6.0). To determine the diagnoses of AMD and RPD, multimodal imaging was performed (Spectralis HRA+OCT, Heidelberg Engineering, Germany; Canon CR6-45NM, Canon, Japan) and graded by a senior retinal specialist (RHG). Optical coherence tomography volume scans were centred on the macula and comprised 49 horizontal B-scans (20° × 20°). The en-face modalities of colour fundus (50°), fundus autofluorescence (30°) and near-infrared imaging (30°) were also centred on the macula. All cases had to have imaging of sufficient quality to be able to grade for AMD stage and RPD status. Beckman classification was used for AMD staging. [91]. Positive RPD phenotyping required at least five RPD lesions seen on two or more B-scans and confirmed on at least one en face modality. For this study, only eyes where RPD made up at least 50% of the total deposits were included in the AMD/RPD+ cases (extensive RPD). Negative RPD phenotyping was defined as < 5 RPD lesions.

### Biopsy processing and fibroblast culture

Skin biopsies were obtained from non-sun-exposed regions using a 3 mm^2^ dermal punch, stored in cold biopsy collection medium (DMEM high glucose, 100 U/mL penicillin and 100 mg/mL streptomycin and 250 ng/L fungizone) and processed on the day of collection. Fibroblasts were expanded, cultured, and banked in DMEM high glucose, 10% fetal bovine serum, L-glutamine, 100 U/mL penicillin and 100 mg/mL streptomycin (all from Thermo Fisher Scientific, USA). All cell lines were mycoplasma-free. Fibroblasts at passage (p) 2 were used for reprogramming.

### Generation, selection, maintenance and quality control of iPSC lines

The reprogramming, maintenance and passaging of iPSCs were performed as described [92]. Patients’ iPSCs were generated by nucleofection (Neon™ Transfection System, Thermo Fisher Scientific) of episomal vectors using Epi5™ Episomal iPSC Reprogramming kit (Thermo Fisher Scientific) in feeder-free and serum-free conditions using TeSR™-E7™medium (Stem Cell Technologies) as we previously described [92]. Pluripotent cells were selected using anti-human TRA-1-60 Microbeads (Miltenyi) [93], maintained onto vitronectin XF™-coated plates (Stem Cell Technologies) in StemFlex™ (Thermo Fisher Scientific), with media changes every 2-3 days and weekly passaging using ReLeSR™ (Stem Cell Technologies). The iPSC lines from AMD/RPD+ (MBE-03537, MBE-03393, MBE-03536,MBE-03579, and MBE-03589), AMD/RPD− (MBE-03556, MBE-03587, MBE-03590, MBE-03408, MBE-03409, TOB-01932) and healthy controls (WAB-00005, WAB-00014, WAB-00025, WAB-00033, WAB-00134, WAB-00546, TOB-01986) were previously generated and characterised in [41,89,94], respectively. Pluripotency was assessed by expression of OCT3/4 (sc-5279, 1:40, Santa Cruz Biotechnology) and TRA-1-60 (MA1-023-PE, 1:100, Thermo Fisher Scientific) by immunocytochemistry with nuclei counterstained using bisBenzimide Hoechst 33342 trihydrochloride (#B2261, Sigma-Aldrich). Virtual karyotyping by CNV analysis, as we previously described [92], using PennCNV and QuantiSNP with default parameter settings. Chromosomal aberrations were deemed to involve at least 20 contiguous SNPs or a genomic region spanning at least 1.5 Megabases.

### Differentiation of iPSCs into RPE cells

RPE cells were generated as described [95]. Briefly, iPSCs were plated on vitronectin (STEMCELL Technologies, 07180) diluted with CellAdhere buffer (STEMCELL Technologies #07183) at a final concentration of 10 µg/mL and hiPSCs were maintained in a 37°C, 5% CO_2_ incubator with daily media changes with Stemflex (Life Technologies, A3349401) until iPSC colony confluency reached 60%. StemFlex medium was replaced with RPE Progenitor Specification Medium, TeSR-E6 (STEMCELL Technologies, 05946) supplemented with N2 (ThermoFisher Scientific, 17502048) with media changes occurring every second day. After 30 days, the medium was switched to RPEM ((MEMα) (Life Technologies, 12561072) supplemented with 5% foetal bovine serum (FBS) (Life Technologies, 26140079), MEM NEAA (Life Technologies, 11140-050, 0.1 mM), N1 supplement (Sigma Aldrich, N6530-5ML, 0.1 mM), 1% L-Glutamine–Penicillin–Streptomycin solution (Sigma Aldrich, G1146-100ML), taurine (Sigma Aldrich, T-0625, 250 µg/mL), hydrocortisone (Sigma Aldrich, H6909, 20 ng/mL) and triiodothyronine (Sigma Aldrich, T-5516, 100 ng/mL)). Cells were subsequently passaged with 0.25% trypsin-EDTA and plated onto growth factor-reduced Matrigel-coated plates (#354230, Corning) at a final concentration of 0.035 mg/cm^2^ to enrich in RPE cells for a final 30 days (90 days total).

### Immunocytochemistry

At Day 60, hiPSC-derived RPE cells were plated onto Matrigel-coated 96-well CellCarrier Ultra plates (PerkinElmer, 6055300; 0.035□mg/cm²). After 30 days in culture with media changes every other day, cells were fixed with 4% paraformaldehyde (10□min, room temperature) and immunostained using standard procedures with the following primary and secondary antibodies: ZO-1 (339100, 10□μg/mL, Life Technologies), PMEL (ab137062, 5□μg/mL, Abcam), RPE65 (ab235950, 10□μg/mL, Abcam), CRALBP (MA1-813, 10□μg/mL, ThermoFisher), AlexaFluor 647 Phalloidin, AlexaFluor 488 goat anti-mouse IgG and AlexaFluor 568 goat anti-rabbit IgG (ThermoFisher Scientific, A22287, A11029, A11011 respectively). Nuclei were counterstained with Hoechst 33342 (Sigma, B2261). Staining specificity was confirmed using appropriate isotype controls. Imaging was performed by full-plate scans (5 fields/well) at 20× magnification on a PerkinElmer Operetta system.

### Transmission electron microscopy

RPE cells were dissociated using 0.25% Trypsin-EDTA (Life Technologies, 25200072), pelleted (5 min, 300□g), and fixed in 2.5% EM-grade glutaraldehyde (Electron Microscopy Sciences, 111-30-8) in 1× sodium cacodylate buffer (Sigma, C0250) for 2□h at room temperature, then stored at 4□°C. After 24□h, cells were washed, resuspended in 0.5% glutaraldehyde, and maintained at 4□°C until shipment. On the day of shipment, cells were pelleted, resuspended in cacodylate buffer, and shipped at ambient temperature to the Institut des Neurosciences de Montpellier (France) for processing and imaging. Upon receipt, cells were immersed in 2.5% glutaraldehyde in PHEM buffer (pH 7.4) overnight at 4□°C, rinsed in PHEM, and post-fixed with 0.5% osmium tetroxide and 0.8% potassium hexacyanoferrate trihydrate for 2□h at room temperature in the dark. Samples were dehydrated through a graded ethanol series (30–100%) and embedded in EmBed 812 using a Leica EM AMW microwave tissue processor. Ultrathin sections (70□nm) were cut using a Leica-Reichert Ultracut E, stained with 1.5% uranyl acetate in 70% ethanol and lead citrate, and imaged using a Tecnai F20 transmission electron microscope at 120□kV (INM, Université de Montpellier, INSERM U1298).

### Transepithelial electrical resistance assay

The electrodes of the epithelial voltohmmeter (EVOM) were first rinsed with 70% ethanol. The EVOM was normalised and a blank measurement was recorded by submerging the electrodes in RPEM. All transepithelial electrical resistance measurements were made under sterile conditions in a cell culture hood, and each sample was measured with the basolateral probe touching the well bottom and with the chopstick electrode at the same angle of approach to avoid significant variability in transepithelial electrical resistance measurements. Net transepithelial electrical resistance measurements were calculated by subtracting the blank value (Matrigel-coated filter without cells) from the experimental value. Each reading was performed in duplicate from two independent experiments. Final resistance-area products (Ω·cm^2^) were obtained by multiplying by the effective growth area.

### Photoreceptor outer segment harvest and assay

Photoreceptor outer segments (POS) were purified from fresh porcine eyes as described previously [96]. Briefly, full retinas were collected under dim red light in a homogenizing solution (20% sucrose, 20 mM tris acetate pH7.2, 2 mM MgCl_2_, 10 mM glucose, 5 mM taurine) and shaken thoroughly. Retinal homogenates were filtered and deposited onto continuous 25-60% sucrose gradients (tris acetate pH7.2, 10 mM glucose, 5 mM taurine). After ultracentrifugation (50 minutes, 25,000 rpm, 4°C), the orange band containing POS was collected and diluted 1:4 in wash solution 1 (20 mM tris acetate pH7.2, 5 mM taurine). POS were then centrifuged (10 minutes, 5,000 rpm, 4°C), washed further in solutions 2 and 3 (10% sucrose, 20 mM tris acetate pH7.2, 5 mM taurine; 10% sucrose, 20 mM Na phosphate pH7.2, 5 mM taurine), with centrifugation steps in between each wash. POS were labeled using 1 mg/mL fluorescein isothiocyanate in DMEM (90 minutes, RT). POS were washed twice in wash solution 3 and once in DMEM. After a last resuspension in DMEM, POS were counted, aliquoted after addition of sucrose to a final concentration of 2.5% and stored at −80°C. Phagocytosis was assessed by flow cytometry following incubation of RPE cells with FITC-labelled POS (FITC-POS). Freshly thawed FITC-POS were diluted into CO_2_-independent medium (ThermoFisher, #18045088) and incubated with RPE cells (200,000 FITC-POS/cm^2^) for 3.5 hours at 37°C. RPE cells were washed twice with filtered PBS^+/+^ (Life Technologies, 14040-133) to remove unbound POS, then RPE were dissociated using trypsin-EDTA (0.25%), phenol red (Life Technologies, 25200072) at 37°C to dislodge cells and remove bound POS from the cell surface. Trypsin was inactivated with RPEM, cells were triturated to generate a single cell suspension containing RPE cells and bound FITC-POS. RPE cells were incubated with DAPI (0.1 µg/mL, Miltenyi, 130-111-570), passed through a 35 µm cell strainer, then analysed on a Becton Dickinson FACS Aria III flow cytometer. Analysis was performed using FCS Express 5 (DeNovo Software). The percentage of RPE cells that internalised FITC-POS was measured from the DAPI-negative gated live cell population. This population was initially gated by size (Forward vs Side scatter) to exclude unbound POS and any cell debris.

### Drusen-like deposit quantification

Confocal imaging was performed using a Zeiss LSM 900 confocal microscope equipped with 2 fluorescence GaAsP, 1 Airyscan detector and transmitted light ESID detector along with 4 diode lasers (405, 488, 561 and 640 nm). The 405, 488, and 561 nm lasers and transmitted light were used in each imaging session. Images were acquired from samples fixed in a 96-well Cell Carrier Ultra Plate in immersion media PBS^-/-^ (Life Technologies, 14190-144) using a 20x/0.8 NA Air Objective. Upon instrument initialization, the plate was calibrated in the Zen 3.2 software and three random points for unbiased imaging were distributed across each well. It was common for the thickness across samples to be highly variable, and it was thus essential that the final Z-stack range was set to accommodate the thickest sample. Automated Z-stack centering was deployed using autofocus during the scan. ZO-1 (excited with 488 nm), APOE (excited with 561 nm), Hoechst (excited with 405 nm) and brightfield (transmitted light) channels were imaged with a Z-stack interval of 0.54 µm over a range of 50 µm (∼100 sections). The output of the semi-automated scan was three 50 μm Z-stacks (.czi files) per well. Imaris (v9.9, Oxford Instruments) was used for processing the Z-stacks acquired during the Zeiss LSM 900 overnight scan. The Zen image files (.czi) were transferred to the Imaris arena and converted to an Imaris file format (.ims) using an Imaris File Converter (v9.9.1). Using the optimised image processing and quantification parameters described below, the software renders a 3D version of the Z-stack and volume-fills each dye. This allowed for quantification of the amount and volume of the drusen-like deposits, measurements that were then exported to .csv file for further statistics. Using Imaris batch analysis, conditions were optimised and individual images were modified in the following workflows: Baseline Subtraction - the background value to be subtracted for each channel was determined using a region selection tool to randomly select three background areas to measure the mean intensity in FIJI. The average of these three measurements was used as the background value with grey values of 6666 and 1111 measured for the APOE (Alexa Fluor 568) channel and ZO-1 (Alexa Fluor 488) respectively. The Baseline Subtraction tool in Imaris was then used to subtract this baseline value from the intensity of every voxel in the image for the APOE and ZO-1 channels. Median Filter - a median filter (3 × 3 × 1) was applied to the Hoechst 33342 channel to reduce background intensity while preserving the edges of the nuclear structure. Surfaces - image segmentation to identify nuclei and drusen-like deposits was performed using the Surfaces tool in Imaris. This tool was used to render a 3D object representing the fluorescent volume in the Z-stack of each channel. The resulting slices were used to aid in visualisation alongside quantitative analysis including spatial and intensity statistics such as deposit count, cell count, and volume of the deposits.

### N-Retinylidene-N-Retinylethanolamine synthesis and treatment

N-Retinylidene-N-Retinylethanolamine (A2E) was synthesised following the protocol in [97]. A2E treatment was carried out in RPE cells cultivated in 96-well Cell Carrier Ultra plates. A2E concentration and treatment schedule (10 µm, every other day, in the dark, for 7 days) were established based on [98].

### RPE cell harvest and single-cell preparation

RPE cells were dissociated using trypsin-EDTA (0.25%) in phenol red (Life Technologies, 25200072) for 5 min at 37°C in a 5% CO_2_ incubator, the plate was tapped to release any non-RPE cells and washed with PBS^-/-^. Trypsin-EDTA (0.25%) in phenol red was added a second time and incubated for 10 min at 37°C in a 5% CO_2_ incubator. The trypsin was inactivated with RPE medium and cells were then harvested into a 15 mL conical tube, as a single-cell suspension. The cell suspension was centrifuged (5 minutes, 300 g, 4°C), then resuspended in 1 mL of 0.1% BSA-PBS. Cells were counted and assessed for viability with trypan blue, and pooled (eight samples maximum) at a concentration of 1000 live cells/μL (1 × 10^6^ cells/mL). The pooled samples were combined with the master mix containing the reverse transcriptase reaction and added to the first well of the third row of the 10x Genomics Chromium Test Chip. Partition oil was loaded into the first well of the first row, and gel beads were added to the first well in the second row. The remaining wells were filled with 50% glycerol and the bottom row was empty. As the chip runs, an emulsion occurs wherein each oil droplet contains one bead and one single cell termed GEM.

### Single cell 3’ RNA-sequencing and pre-processing of transcriptome data

Single cell RNA-sequencing libraries were prepped using the Chromium Single Cell 3’ Library & Gel bead Kit for 10X Genomics. Libraries were sequenced on Illumina NovaSeq 6000. Raw base calls (BCL files) from the sequencer were demultiplexed into FASTQ format using the *bcl2fastq* software (https://sapac.support.illumina.com/sequencing/sequencing_software/bcl2fastq-conversion-software.html). Read quality control, alignment, cell and UMI counting was performed using Cell Ranger *count* pipeline from 10X Genomics (v6.0.2 was run on the first batch and v7.1.0 on the remaining three batches) with the default target cell count at 20,000 and Homo Sapiens GRCh38 (GRCh38-2020-A) as the reference. Quality control filters were tailored to each library to account for differences in sequencing depth. The aim was to remove the lower and upper outlier barcodes. Additionally, barcodes that could not be confidently assigned to a donor or were deemed a doublet were also excluded (see the *Demultiplexing of cell pools into individual donors* section for details). The mitochondrial gene expression threshold was set to 30% of the total transcripts. Normalisation and scaling were performed using *Seurat’s* SCTransform method with ribosomal gene content percentage regressed out. All libraries were then merged and integrated using *Harmony* R package (version 1.2.0) with the sequencing pool as a covariate. Sample variables like sex and age were tested and were not found to cause batch effects.

### SNP genotyping and imputation

DNA was extracted from cell pellets using QIAamp DNA Mini Kit (QIAGEN, 51306) as per the manufacturer’s instructions. DNA concentration was determined with a SimpliNano spectrophotometer (GE Life Sciences), and DNA was diluted to a final concentration of 10-15 ng/µl. Samples were genotyped on the UK Biobank Axiom™ Arrays (Ramaciotti Centre for Genomics, Sydney, Australia). Pre-processing of the raw CEL files and VCF generation was done using Axiom Analysis Suite (version 5.3.0.45) from Thermo Fisher Scientific. Genotype imputation was run using the SNP_imputation_1000g_hg38 pipeline using the 1000G Project as the reference (https://github.com/powellgenomicslab/SNP_imputation_1000g_hg38/tree/master). Imputation quality filters were set to minor allele frequency (MAF) of >10%, R^2^ score of 0.8 and fraction of missing genotypes of 0.5, resulting in 4,063,692 SNPs.

### Demultiplexing of cell pools into individual donors

Demultiplexing and doublet detection for the libraries was run with *Demuxafy* (version 3.0.0) using the SNP genotyping data as reference (https://demultiplexing-doublet-detecting-docs.readthedocs.io/en/latest/Background.html). The softwares chosen for demultiplexing were *demuxlet*, *demuxalot* and *souporcell*, while *scds* and *scDblFinder* were used for doublet detection. The parameters for all methods were set to the defaults in *Demuxafy*. The results were consolidated using the “MajoritySinglet” approach in combine_results.R of *Demuxafy*. The final assignments were incorporated into the Seurat objects.

### Classification of cell subpopulations

The first 20 principal components from Harmony embedding were chosen for the Uniform Manifold Approximation and Projection (UMAP) visualisation. The optimal clustering resolution was deduced as 0.3 using the gap statistic method from scBubbleTree which resulted in 14 clusters. Canonical marker genes associated with RPE and proliferative functions were examined for each cluster, as we previously described [41]. As a first pass, we performed label transfer using the previously described RPE cells as reference [38]. Additionally, markers for each of the clusters were also investigated using Seurat’s *FindMarkers* function. Clusters that undoubtedly demonstrated proliferative markers were labelled as RPE progenitors. The remaining clusters showed varying levels of RPE functions and were deduced as either early RPE or RPE.

### Differential expression analysis

Since the main focus of this study was understanding the variance within disease phenotypes rather than specific states of maturity, we combined all RPE subpopulations (early and mature RPE) into a single group and interrogated differences between the two disease phenotypes within this combined group. This helped increase the power of the calculations. Moreover, in some of the smaller RPE populations, not all donors were represented. The analysis was performed using the MAST (v1.28.0) implementation within the Seurat package using log-transformed normalised counts from SCTransform. Covariates included as latent variables include total UMI count, sequencing pool, sex and age. Thresholds for significant results were set to | Average Fold Change | > 0.25 and adjusted *p* value < 0.05.

### Gene ontology and disease ontology over-representation analysis

For each disease phenotype, gene sets for over-representation analysis were prepared using the significant hits from differential expression analysis. Over-representation analysis was performed with the *clusterProfiler* (v4.10.1) package [50,99] to identify enriched pathways and processes using three resources: Gene Ontology, Disease Ontology (DOSE v3.28.2) and KEGG pathways.

### Mapping of expression and protein QTL

To explore any association between genetic variation and gene expression in the context of disease states, we performed eQTL analysis using a pseudobulk approach with *matrixEQTL* (v2.3) (https://github.com/andreyshabalin/MatrixEQTL). For each population, an aggregated donor-gene matrix was generated by running quantile normalisation on the mean values of each gene. The analysis was run using an additive linear model on all non-zero genes and 4,063,692 SNPs. For each gene, SNPs located within a 1MB window from start and end of the gene were tested. Covariates specified were age, sex, top five genotype principal components and disease state. Significant eQTLs were filtered with the following thresholds: false discovery rate < 0.05 and homozygous alternate allele frequency of at least 5. Within the subset of significant eQTLs, to test whether the effect of the eQTLs differed between the disease states, we added a Genotype:Phenotype interaction term to the model and filtered by P value < 0.05 of the interaction term.

pQTL analysis was run on bulk-level protein abundance measurements from 69 RPE cell lines. The abundance matrix was normalised using rank-based inverse transformation. Genotype data of the RPE cell lines gave 4,078,773 SNPs (MAF > 10%) after QC. Covariates incorporated into the model were age, sex, top 10 genotype principal components and disease state. Association test for each protein was run with *QTLtools cis* permutation test (v1.3.1) [100] using SNPs within a 1MB window. FDR values were calculated on the variant-level adjusted empirical P-values and filtered by <0.05 for significant pQTLs.

### Transcriptome wide association study

TWAS analysis was performed with the FUSION package [101] with the following inputs: a) gene expression weights calculated with *FUSION.compute_weights.R* using matrixEQTL summary statistics with “blup”, “lasso”, “top1” and “lasso” models and, b) RPD−specific GWAS summary statistics [29]. The publicly available 1000 Genomes LD reference file (https://github.com/gabraham/plink2R/archive/master.zip) was used since we did not have enough samples to create our own. FDR values were calculated on the association P values for each cell type then filtered by 10% for significance.

### Preparation of protein samples

RPE cell cultures were lysed in RIPA buffer containing phosphatase and protease inhibitors, then sonicated (40 Hz, 2 pulses of 15 s each) using a probe sonicator. Insoluble debris was removed by centrifugation at 14,000 rpm for 15 minutes at 4°C. Protein concentrations were measured using the bicinchoninic acid protein assay kit (Thermo Scientific). Extracted proteins (50 μg) were adjusted to 50 μL in 1× S-Trap lysis buffer and reduced/alkylated with TCEP and MMTS. The S-Trap midi protocol (Protifi) was performed. Phosphoric acid was added to the SDS lysate (final concentration 1.2%), followed by the addition of 350 μL S-Trap binding buffer (90% aqueous methanol with 100 mM TEAB). The mixture was loaded onto an S-Trap column, washed, and digested with trypsin at a 10:1 protein ratio in 50 mM TEAB. After overnight incubation at 37°C, peptides were eluted in three stages, pooled, and concentrated using a Speed-Vac to near dryness (∼5 μL remaining).

### Tandem mass tag labelling and peptide fractionation

Proteome profiling was conducted on a Tandem Mass Tag (TMT) platform with six independent 16-plex TMT experiments. Dried peptides were resuspended in 100 mM HEPES buffer (pH 8.2), and concentrations were determined with the MicroBCA kit. Peptides from each sample (35 μg) were labelled with 0.2 mg TMT reagent per tube for 1 hour at room temperature with vortexing. Residual TMT reagent was quenched with 8 μL of 5% hydroxylamine. For each 16-plex experiment, labelled samples were combined and dried by vacuum centrifugation. The peptides were cleaned using a C18 column (Sep-pak, Waters) before undergoing high-pH reversed-phase fractionation on an Agilent 1260 HPLC system. Peptides were separated using a 55-minute gradient from 3-30% acetonitrile in 5 mM ammonia (pH 10.5) at a flow rate of 0.3 mL/min. The collected 96 fractions were consolidated into eight final fractions. These were dried, resuspended in 1% formic acid, and desalted using SDB-RPS stage tips.

### Liquid chromatography and tandem mass spectrometry

Mass spectrometric data were acquired using an Orbitrap Eclipse mass spectrometer connected to a Vanquish Neo UHPLC system. Approximately 1 µg of peptide was separated on a 100 µm capillary column packed with 35 cm of Accucore 150 resin (2.6 μm, 150Å; ThermoFisher Scientific) at a flow rate of 350 nL/min. The scan sequence started with an MS1 spectrum (Orbitrap analysis, resolution set to 60,000, mass range 350-1350 Th, automatic gain control (AGC) target set to 100%, maximum injection time set to “auto”. Data acquisition lasted 75 minutes per fraction. The high-resolution MS2 (hrMS2) stage involved fragmentation via higher-energy collisional dissociation (HCD) at a normalized collision energy of 36%. Analysis was performed using the Orbitrap (AGC target 200%, maximum injection time 86 ms, isolation window 0.5 Th, resolution 30,000, with TurboTMT enabled). The FAIMSpro interface was used, with a dispersion voltage (DV) of 5,000V and compensation voltages (CVs) set at +30V, −50V, and −70V. The TopSpeed parameter was set to 1 second per CV.

### Proteomic data analysis

Spectra were converted to mzXML format using MSconvert [102]. Database searches were conducted using all entries from the homo sapiens UniProt reference database (downloaded: June 2024) and a reversed concatenated sequence database. Searches were performed using a 50-ppm precursor ion tolerance for protein profiling and a product ion tolerance of 0.03 Da, maximizing sensitivity for Comet searches and linear discriminant analysis [103,104]. Static modifications included TMTpro labels on lysine residues and peptide N-termini (+304.207 Da), along with carbamidomethylation of cysteine residues (+57.021 Da). Oxidation of methionine residues (+15.995 Da) was set as a variable modification. Peptide-spectrum matches (PSMs) were adjusted to a 1% false discovery rate [105,106]. PSM filtering utilized linear discriminant analysis [104,106], and proteins were assembled to a final protein-level false discovery rate of 1% [106]. Protein quantification was based on summing reporter ion counts from all matching PSMs [107]. Reporter ion intensities were corrected for isotopic impurities according to manufacturer specifications. The signal-to-noise (S/N) measurements for peptides assigned to each protein were summed, and these values were normalized to ensure equal protein loading across channels. Each protein abundance was then scaled such that the summed signal-to-noise ratio for that protein across all channels equaled 100, resulting in a relative abundance (RA) measurement. DEPs were identified through Student’s *t*-tests, specifically comparing the ratios between both groups. The overall fold changes were calculated as geometric means of the corresponding ratios. For a protein to be considered differentially expressed, we followed the following two criteria: a ratio fold change exceeding 1.2 for upregulated or falling below 0.83 for downregulated expression, and a *p* value cutoff (*t*-test *p*□<□0.05). Investigation of protein–protein interactions and functional enrichment gene ontology (GO) analysis of differentially expressed proteins (DEPs) were performed with the online STRING database version 12.0119. STRING analysis on the 715 DEPs within the proteomics dataset, generated a network of interactions (based on both evidence of functional and physical interactions). The top hits were selected based on the minimum required interaction score of 0.9 (highest confidence), and the full STRING network was deployed to visualise network maps and interactors. Network lines represent the protein interaction score, which was set at a minimum high confidence (0.7) for visualisation. Active interaction sources were based on text mining, experiments, databases, coexpression, neighbourhood, gene fusion, and co-occurrence data. The network analysis uses a strength score of each STRING network, which is the ratio between the number of proteins in the specified network that are annotated with a term, and the number of proteins that STRING expects to be annotated with this term in a random network of the same size (Log10(observed/expected)=strength).

## Data availability

All data supporting the findings of this study are available within the paper and its Supplementary Information. Sequencing and processed transcriptomic data are in the process of being deposited to a public repository and will be made openly available upon publication. Proteomics raw data and search results are being deposited to the ProteomeXchange Consortium via the PRIDE [108] partner repository and will be released upon publication.

## Supplementary Information

Supplementary Data. All datasets

Supplementary Info. Additional figures, table, movie.

## Acknowledgments

We thank all participants who donated skin biopsies. The authors acknowledge the facilities and the scientific and technical assistance of the iPSC reprogramming facilities, the Stem Cell Disease Modelling laboratory, the University of Melbourne, which is supported by Phenomics Australia (PA) through funding from the Australian Government’s National Collaborative Research Infrastructure Strategy program. Microscopy was performed at the Biological Optical Microscopy Platform Facility at The University of Melbourne. RNA sequencing was performed at the Garvan-Weizmann Centre for Cellular Genomics and genotyping at the Ramaciotti Centre for Genomics.

## Authors’ Contributions

Conceptualisation: J.C.H., K.K.S., M.D., A.S., A.W.H., R.C.H., J.E.P., A.P.; Methodology: J.C.H., K.K.S., M.D., A.S., C.J.A., G.E.L., M.M, A.W.H., R.C.H., J.E.P., A.P.; Investigation: J.C.H., K.K.S., M.D., A.S., H.H.L., H.K., J.M., T.A., Y.H., E.F.N., C.C., G.M., S.M., P.T.; Data analysis: J.C.H., K.K.S., M.D., A.S., M.M., A.B., D.P., J.E.P., A.P.; Writing original draft: J.C.H., K.K.S., M.D., A.S., A.P.; Writing review & editing, all authors; Funding Acquisition: B.R.E.A., E.L.F., Z.W., M.B., R.H.G, A.P.; Supervision and project administration: A.P.

## Funding

This research was supported by a National Health and Medical Research Council (NHMRC) Synergy grant 1181010 (ELF, ZW, MB, BREA, RHG, AP, co-led by the RPD Consortium principle investigators and collaborators, Appendix 1), an NHMRC Early Career Fellowship (GNT1157776, BREA), NHMRC Investigator Grants (GNT2008382, ZW; GNT1195236, MB; AWH; GNT1194667, RHG; JEP) an NHMRC Senior Research Fellowship (1154389, AP), a Dame Kate Campbell Fellowship (AP), the Australian Research Council Training Centre for Personalised Therapeutics Technologies (IC170100016, JCH, AP), a IHU FOReSIGHT from Agence Nationale de la Recherche (ANR-18-IAHU-0001, EFN), Centre National de la Recherche Scientifique (EFN, Tenure position), the University of Melbourne, the NHMRC Independent Research Institute Infrastructure Support Scheme (IRIISS) and Operational Infrastructure Support from the Victorian Government.

## Materials & Correspondence

Requests for materials and correspondence should be addressed to Alice Pébay.

## Competing Interests

A.W.H., J.E.P., and A.P. are directors and shareholders of CellTellus Pty Ltd.

---

**Figure S1.**
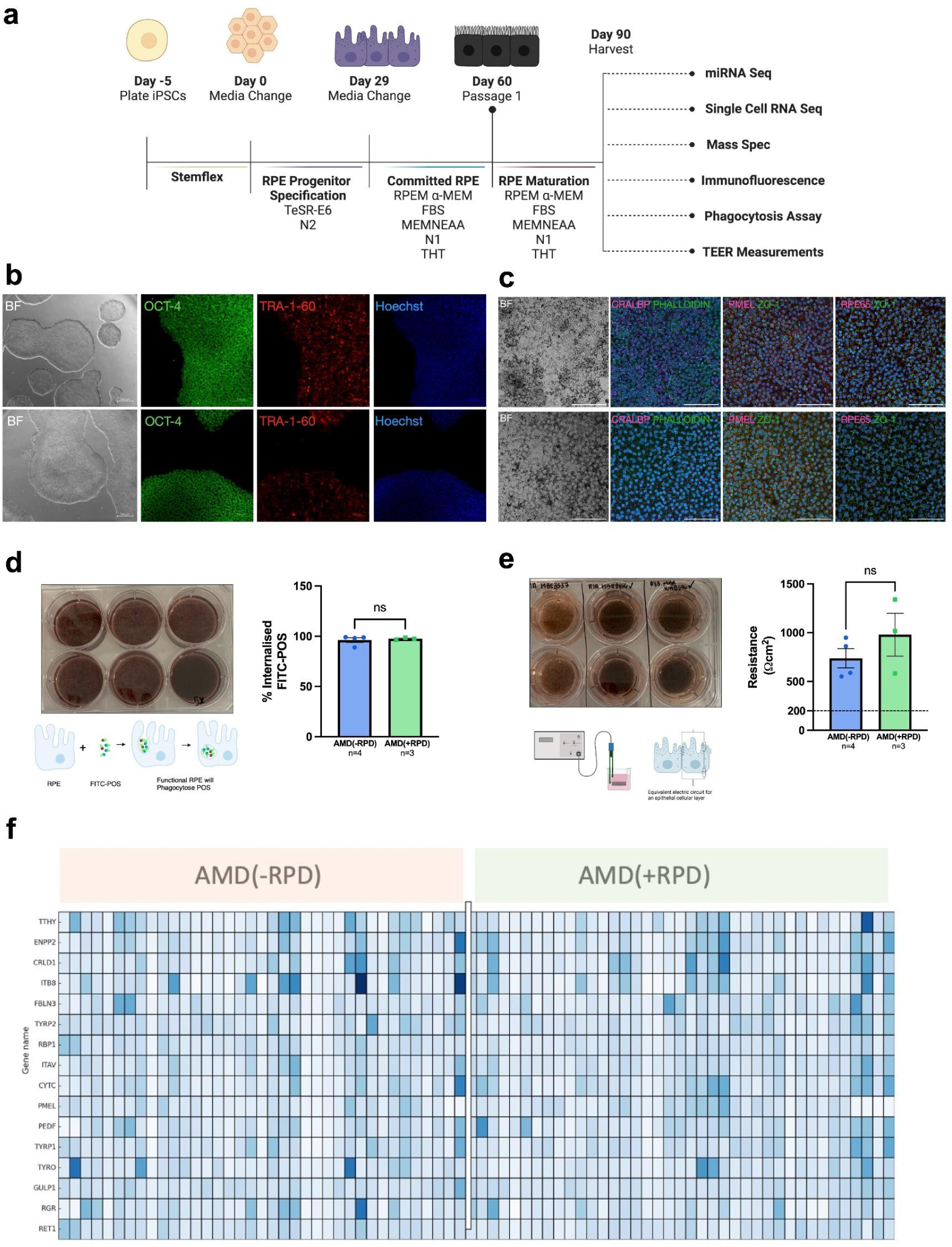
Quality control. (**a**) Schematic overview of the experimental workflow (created with BioRender.com). (**b**) Quality control of generated iPSC lines showing representative brightfield (BF) and immunofluorescence images for OCT-4, TRA-1-60, and Hoechst 33342 in a AMD/RPD− line (MBE-03556, top) and an AMD/RPD+ line (MBE-01986, bottom). Scale bars: 500 µm (BF), 100 µm (fluorescence). Images are representative of all lines. (**c**) Characterisation of iPSC-derived RPE cells from the same two lines, showing BF and immunostaining for phalloidin (magenta) and CRALBP (green); PMEL (magenta) and ZO-1 (green); RPE65 (magenta) and ZO-1 (green). Scale bars: 100 µm. (**d**) Phagocytosis assay. Representative pigmented RPE cells and schematic (BioRender) showing uptake of FITC-labelled photoreceptor outer segments (POS). Functional assay results for AMD/RPD− (MBE-03556, MBE-03556, MBE-03556, MBE-03556) and AMD/RPD+ (TOB-01986, TOB-01986, TOB-01986) lines, presented as mean % engulfment ± SEM from technical triplicates of each unique iPSC-derived RPE line. Significance was assessed by t-test (p < 0.05). (**e**) Transepithelial electrical resistance (TEER) assay. Representative pigmented RPE cells on transwell inserts and schematic (BioRender) of the TEER setup. Resistance values are shown for AMD/RPD− (MBE-03556, MBE-03556, MBE-03556, MBE-03556) and AMD/RPD+ (TOB-01986, TOB-01986, TOB-01986) lines. Each dot represents the average of technical triplicates for each iPSC-derived line. Data are shown as mean ± SEM, with statistical significance determined by *t*-test (p < 0.05). (**f**) Heatmap of RPE canonical marker expression from mass spectrometry across all AMD/RPD− and AMD/RPD+ lines. Each rectangle represents one line; colour intensity reflects raw abundance values from minimal (light blue) to maximal (dark blue).

**Figure S2.**
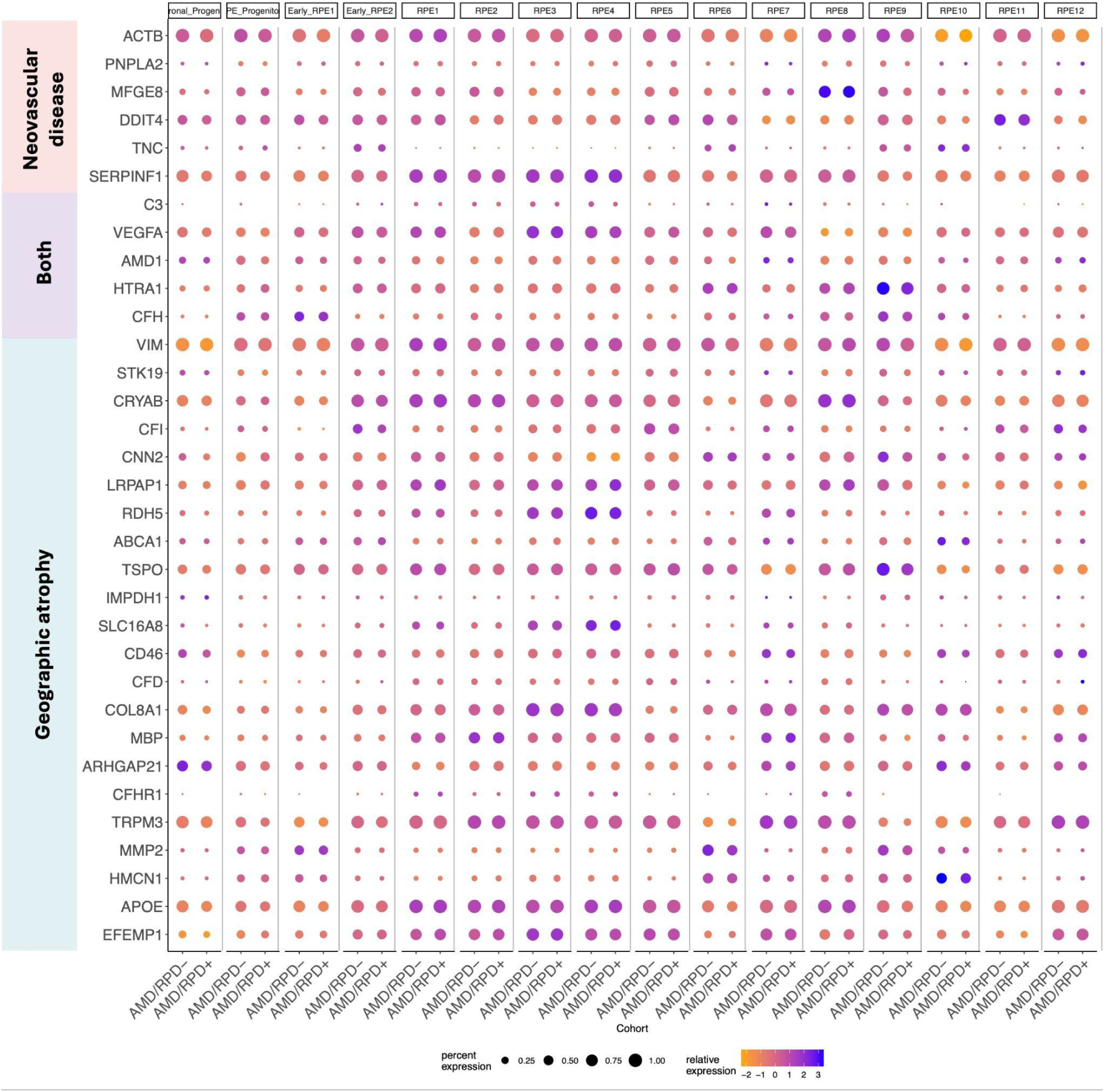
Genes associated with cell subpopulations and their expression in AMD/RPD− and AMD/RPD+ cells. Dotplot representation of single-cell expression profiles for genes linked to AMD. Plots show scaled average expression (z-scores; color scale) and the percentage of cells within each cluster expressing the gene (dot size).

**Figure S3.**
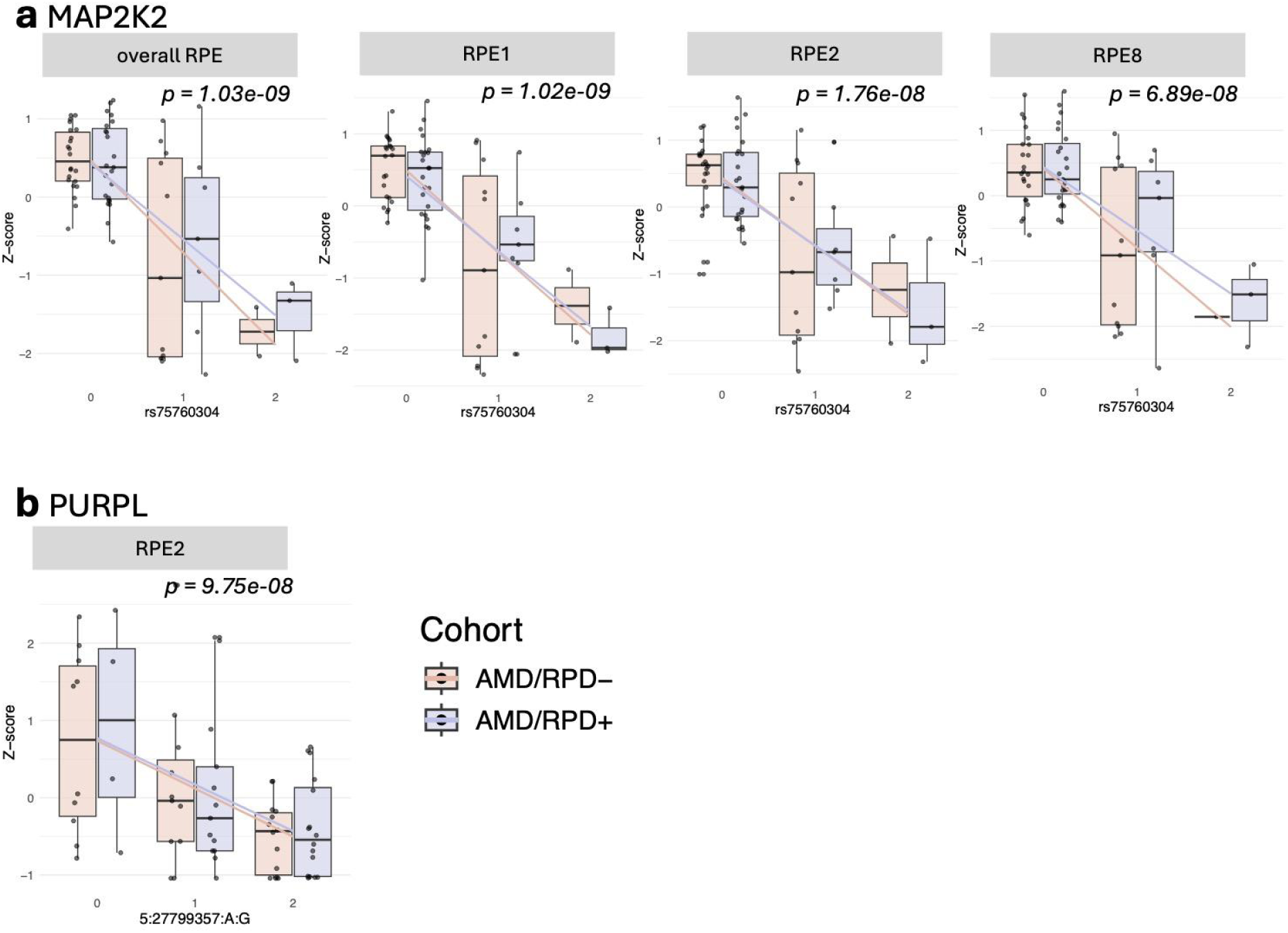
Examples of eQTLs in RPE subpopulations. (**a**) Expression of *MAP2K2* stratified by rs75760304 genotype in overall RPE and in individual subpopulations (RPE1, RPE2, RPE8), shown separately for AMD/RPD− and AMD/RPD+ cohorts. (**b**) Expression of *PURPL* stratified by 5:27799357:A:G genotype in RPE2, shown separately for AMD/RPD− and AMD/RPD+ cohorts. Expression values are represented as z-scores.

**Figure S4:**
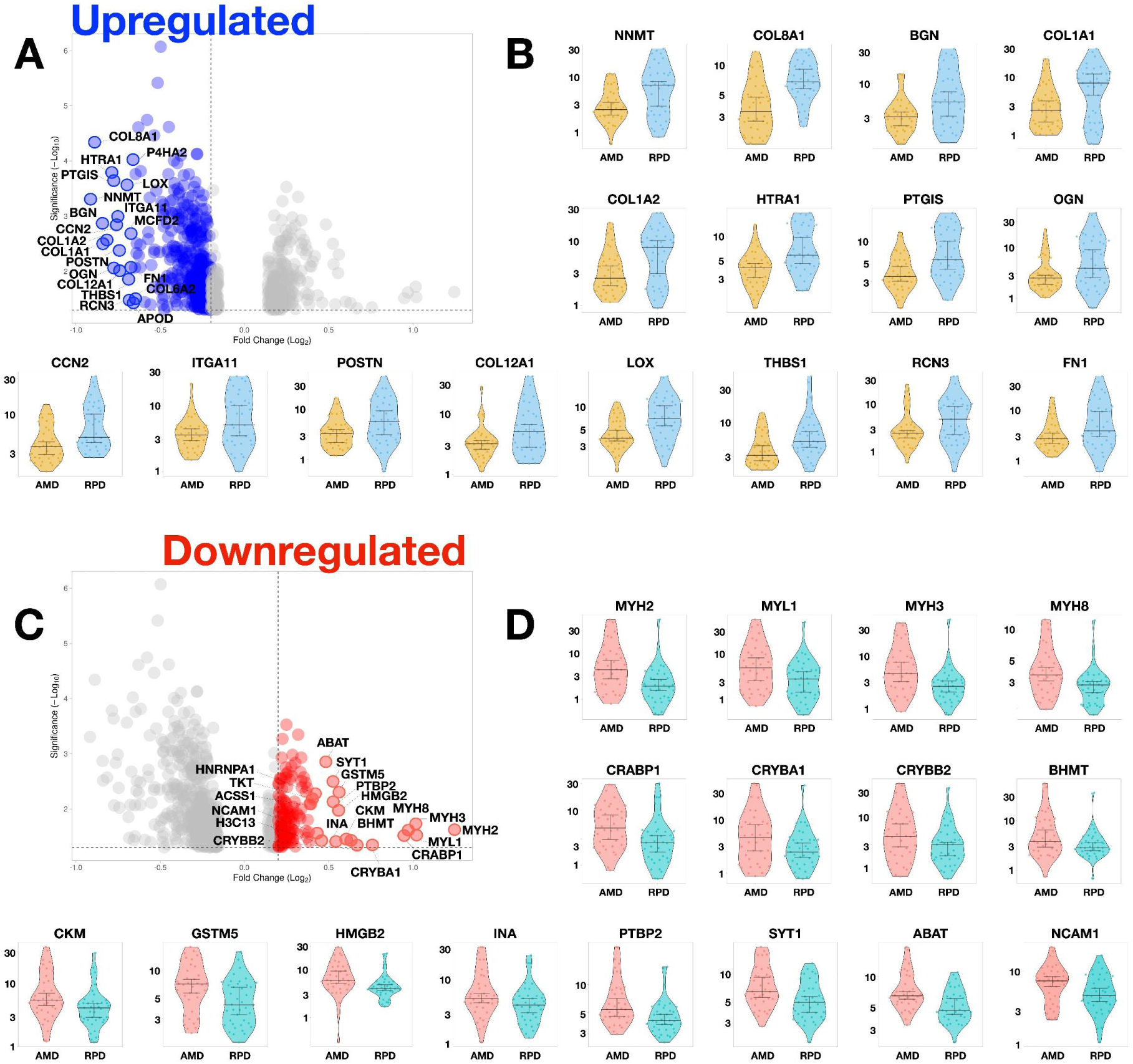
Top differentially expressed proteins between AMD (AMD/RPD−) and RPD (AMD/RPD+) RPE cells. Violin plots showing the top 16 proteins most significantly up- or down-regulated in AMD/RPD+ relative to AMD/RPD− RPE cells based on log₂-normalised TMT proteomic data. Each violin represents the distribution of protein abundance.

**Figure S5:**
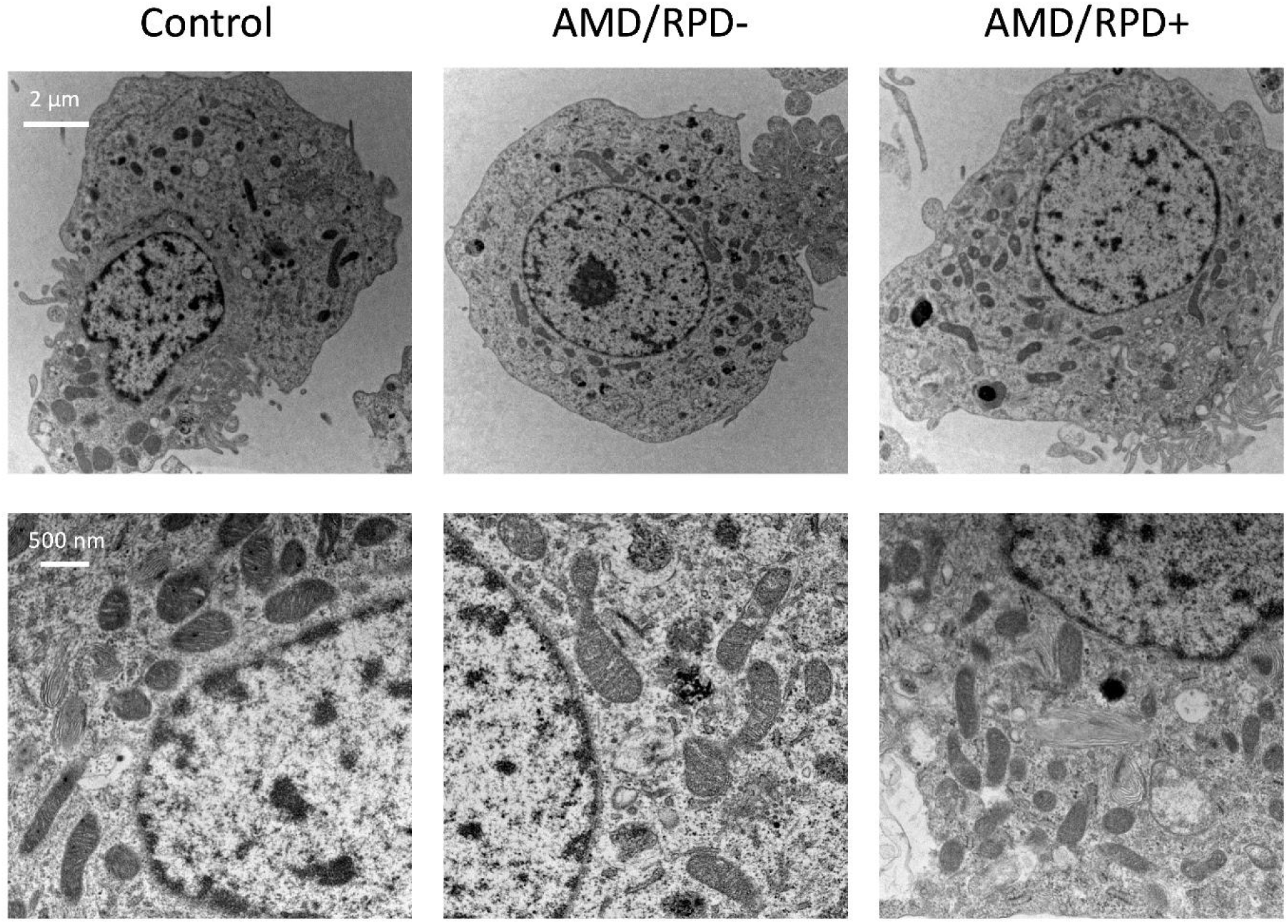
Transmission electron microscopy of iPSC-derived RPE cells. Representative images of RPE cells derived from control, AMD/RPD−, and AMD/RPD+ donors. Top row: low-magnification views showing overall cell morphology, including nuclei, cytoplasmic organisation, and organelles (scale bar: 2 µm). Bottom row: higher-magnification views highlighting mitochondrial ultrastructure and cytoplasmic detail (scale bar: 500 nm). No consistent ultrastructural differences were observed between cohorts.

**Table S1.** Full details of significant pQTLs. Complete information including genomic coordinates and alleles.

| Uniprot ID | Gene | Chr | Pos from | Pos to | Strand | SNP | SNP ID | Slope (beta) | Adj Beta P value | FDR |
| --- | --- | --- | --- | --- | --- | --- | --- | --- | --- | --- |
| Q6YP21 | KYAT3 | 1 | 88935774 | 88936245 | - | 1:88974992:G:A | rs35594043 | -0.784461 | 4.07702e-05 | 0.04351811 |
| Q9Y6B7 | AP4B1 | 1 | 113894195 | 113895492 | - | 1:113905292:G:T | rs971173 | -0.836572 | 1.84237e-05 | 0.03651718 |
| Q8N2H3 | PYROXD2 | 10 | 98383569 | 98383868 | - | 10:98400642:T:C | rs6584194 | 1.14271 | 3.92864e-08 | 0.00020967 |
| Q86SX6 | GLRX5 | 14 | 95535051 | 95535384 | + | 14:95544087:G:A | rs11628901 | -1.16739 | 3.92053e-05 | 0.04351811 |
| P15559 | NQO1 | 16 | 69709402 | 69711281 | - | 16:69687061:G:A | rs3169315 | -0.876558 | 2.05268e-05 | 0.03651718 |

## Appendix

## Appendix 1 RPD consortium contributors #

### Principal Investigators

Robyn Guymer, Erica Fletcher, Melanie Bahlo, Alice Pébay, Brendan Ansell and Zhichao Wu.

### Project manager

Carla Abbott

### Genetic cohorts

Elvira Agron, Lebriz Altay, Ella Arnon, Louis Arnould, Mary Attia, Bjorn Bakker, Konstantinos Balaskas, Melinda Cain, Emily Caruso, Usha Chakravarthy, Jason Charng, Attiqa Chaudhary, Fred K Chen, Shang-Chih Chen, Emily Chew, Sae Cho, Itay Chowers, Linda Clarke, Audrey Cougnard-Grégoire, Catherine Creuzot Garcher, Catherine Cukras, Stéphanie Debette, Cécile Delcourt, Marie-Noëlle Delyfer, Anneke I den Hollander, Layal El Wazan, Sarah Elbaz-Hayun, Sascha Fauser, Daniela Ferrara, Robert P Finger, Pierre-Henry Gabrielle, Javier Gayan, Erin Gee, Emily Glover, Kai Lyn Goh, Michelle Grunin, Rachael Heath Jeffery, Catherine Helmer, Wilson Heriot, Claire Hill, Lauren AB Hodgson, Ruth Hogg, Frank G Holz, Carel Hoyng, Amy Kalantary, Haya Kashtan, Pearse A Keane, Frank Kee, Tiarnan Keenan, Samer Khateb, Meme Ko, Jean-François Korobelnik, Adi Kremer, Himeesh Kumar, Eleonora M Lad, Mélanie Le Goff, Yara Lechanteur, Jackson Lee, Rivkah Lender, Jaime Levi, Sandra Liakopoulos, Timing Liu, Ulrich FO Luhmann, Chi D Luu, Matthias M Mauschitz, Amy J McKnight, Samuel McLenachan, Aniket Mishra, Ismail Moghul, Tunde Peto, Nikolas Pontikos, Anna Rautanen, Batya Rinsky, Danial Roshandel, Danuta M Sampson, Sean Santiago, Tina Schick, Cédric Schweitzer, Merav Shiryon, Michal Shpigel, Yahel Shwartz, Vasilena Sitnilska, Laura Smyth, Amy Stockwell, Jan H Terheyden, Nikita Thomas, Liran Tiosano, Adnan Tufail, Claire Weber, Brian L Yaspan, MACUSTAR Consortium, NICOLA Consortium.

### Other researchers

Yelena Bagdasarova, Roberto Bonelli, Steven Clarke, Maciej Daniszewski, Coen De Vente, Samaneh Farashi, Una Greferath, Satya Gunnam, Jenna Hall, Jiru Han, Marco Herold, Alex Hewitt, Victoria E Jackson, Aaron Lee, Helena Liang, Grace Lidgerwood, Jessica Ma, Luz D Orozco, Joseph Powell, Matt Rutar, Clarisa Sánchez, Roy Schwartz, Liam W Scott, Manisha Shah, Kaylene Simpson, Scott Song, Anand Swaroop, Kirstan Vessey.

